# DYNAMIC CONFORMATIONS IN THE ACTIVATION OF PINK1 KINASE

**DOI:** 10.64898/2026.08.19.745768

**Authors:** Huiqin Xu, Xinyang Liu, Yating Jiang, Yanfeng Zhang, Junrong Xue, Shujuan Wang, Yan Du, Shuaijiabin Chen, Chen Chen, Xinrong Yu, Bingxuan Li, Sen-Fang Sui, Xiaohong Qin, Li-Zhi Mi, Zheng Liu

**Author notes:** Zheng Liu, Li-Zhi Mi, Xiaohong Qin, Sen-Fang Sui, **Email:**. Equal contribution (Huiqin Xu, Xinyang Liu, Yating Jiang, Yanfeng Zhang).

## Abstract

Under mitochondrial stress, the mitochondrial kinase PINK1 is activated to phosphorylate ubiquitin, which in turn recruits the E3 ligase Parkin, thereby initiating clearance of damaged mitochondria. Dysregulation of the PINK1–Parkin signaling pathway is associated with early-onset autosomal recessive Parkinson’s disease. However, the mechanisms governing PINK1 conformational dynamics and activation in response to mitochondrial stress remain poorly understood. Here, we report that *Tribolium castaneum* PINK1 (*Tc*PINK1) forms specific symmetric dimers via interactions between the kinase N-lobe and the regulatory C-terminal domain. These dimers undergo dynamic conformational transitions across distinct states, including an autoinhibited state, an inactive intermediate state, a primed-activation state and an active state. Moreover, *Tc*PINK1 undergoes lateral trans-phosphorylation through dimer–dimer interactions. Notably, symmetric dimer formation is essential for Parkin recruitment during mitophagy. These findings provide a framework for understanding the conformational plasticity and allosteric regulation of PINK1 in mitophagy.

## Introduction

PINK1 (Phosphatase and Tensin Homolog (PTEN)-induced putative kinase 1, PARK6), a mitochondrial kinase, serves as a key regulator of mitochondrial autophagy (mitophagy), motility, and calcium (Ca²⁺) homeostasis (Gandhi, Wood-Kaczmar et al. 2009, Narendra, Jin et al. 2010, Vives-Bauza, Zhou et al. 2010, Wang, Winter et al. 2011). Dysregulation of PINK1 functions due to pathological mutations interferes with the clearance of damaged mitochondria, impairs mitochondrial integrity, and leads to the degeneration of dopaminergic neurons, which is associated with early-onset autosomal recessive Parkinson’s disease (Valente, Abou-Sleiman et al. 2004, Clark, Dodson et al. 2006, Park, Lee et al. 2006, Yang, Gehrke et al. 2006, Dagda, Cherra et al. 2009). Therefore, the activity of PINK1 must be tightly regulated.

Under normal conditions, PINK1 is imported into mitochondria via the translocase of the outer mitochondrial membrane (TOM) complex, cleaved by Presenilin-associated rhomboid-like protein (PARL), and subsequently transported back and degraded to maintain mitochondrial homeostasis. In response to stress, PINK1 importation is stalled so that it is accumulated on the outer mitochondrial membrane, where it is activated through transphosphorylation and/or dimerization, enabling it to recruit Parkin via ubiquitin phosphorylation and to initiate mitophagy (Lazarou, Jin et al. 2012, Okatsu, Uno et al. 2013). These functional transitions inevitably require PINK1 to adopt different conformations and assemblies.

Early structural studies have shed light on PINK1’s conformation, substrate recognition, ATP binding, transphosphorylation, and importation (Kumar, Tamjar et al. 2017, Schubert, Gladkova et al. 2017, Okatsu, Sato et al. 2018, Gan, Callegari et al. 2022, Rasool, Veyron et al. 2022, Callegari, Kirk et al. 2025). However, gaps still remain in understanding the structural basis for PINK1’s activity regulation. Intriguingly, many early structural studies showed PINK1 adopting similar active conformations under varying conditions, despite some kinases being extensively engineered with diminished phosphorylation activity (Kumar, Tamjar et al. 2017, Okatsu, Sato et al. 2018, Gan, Callegari et al. 2022, Rasool, Veyron et al. 2022). This raises questions about the how PINK1 conformations and activities are regulated.

Recent studies revealed that insect and human PINK1s can form face-to-face dimers, either within dodecameric complex or in association with TOM complex (Gan, Callegari et al. 2022, Rasool, Veyron et al. 2022, Callegari, Kirk et al. 2025). However, the two kinases within these dimers are mutually blocking each other from access to additional substrates, including ubiquitin, Parkin, and PINK1, complicating the understanding of how they can facilitate transphosphorylation of their substrates and recruit Parkin to damaged mitochondria.

In early structural studies, the term “activation” is used to describe PINK1’s transition from a resting to a stimulated state, yet its definition and assessment criteria are inadequately defined. Based on published reports, we believe that a complete activation process of PINK1 should include: 1) autoinhibition in the absence of stress; 2) association with TOM complex in response to membrane depolarization; 3) formation of different complexes in the transphosphorylation of PINK1, ubiquitin, and Parkin; 4) stabilization in an activated state; and 5) recruitment of Parkin for mitophagy. Thus, the criteria for PINK1 activation should encompass: 1) PINK1’s conformation; 2) its autophosphorylation activity; 3) its activity in ubiquitin transphosphorylation; and 4) its ability to recruit Parkin in mitophagy. Defining these criteria is crucial for elucidating the structural determinants for PINK1’s activity regulation.

To this end, we developed a fluorescent-based assay to measure PINK1 transphosphorylation and autophosphorylation activities, determined cryo-EM structures of *Tc*PINK1 kinases, and analyzed the effects of PINK1 mutations on Parkin recruitment and mitophagy. Our findings reveal that *Tc*PINK1 form back-to-back dimers through interactions between the kinase N-lobe and C-terminal regulatory domain. These dimers dynamically interconvert among autoinhibited, prime-activated, inactive transitional, and active states, engaging in the Parkin recruitment and mitophagy regulation. Furthermore, *Tc*PINK1 can trans-phosphorylate each other through dimer- dimer interactions. Collectively, these results elucidate the molecular basis for PINK1’s conformational dynamics and assembly in mediating Parkin recruitment in mitophagy.

## Results

### The Dimeric, Activating Mutant, *Tc*PINK1^L552R^, Is Dynamically Populated in Two Distinct Conformations

From the multiple sequence alignment, we found that the N-terminal fragment of PINK1 αA helix, from the 121^st^ to 134^th^ a.a. of *Tc*PINK1, is highly variable in length and amino acid composition (Fig. S1). The corresponding fragment from *Gekko japonicus* PINK1 has 11 a.a., including 2 Prolines, while the fragment from *h*PINK1 has 18 a.a.. We therefore truncated the N-terminal fragment of *Tc*PINK1 up to the 134^th^ a.a., developed a high-sensitive fluorescent-based kinase activity assay, and identified that L552R mutation could stabilize *Tc*PINK1 in an active, dimeric state (Fig. S2, S3).

To further define the molecular basis for the dimerization and activation of *Tc*PINK1, we determined the Cryo-EM structures of this mutant (hereafter referred to as *Tc*PINK1^L552R^) at resolutions of 2.8 to 3.35 Å (Fig. S4, Table S1). Remarkably, this mutant is dynamically populated in two distinct conformations: one representing an autoinhibited state (Fig. 1) and the other primed for activation (Fig. S5).

**Figure 1.**
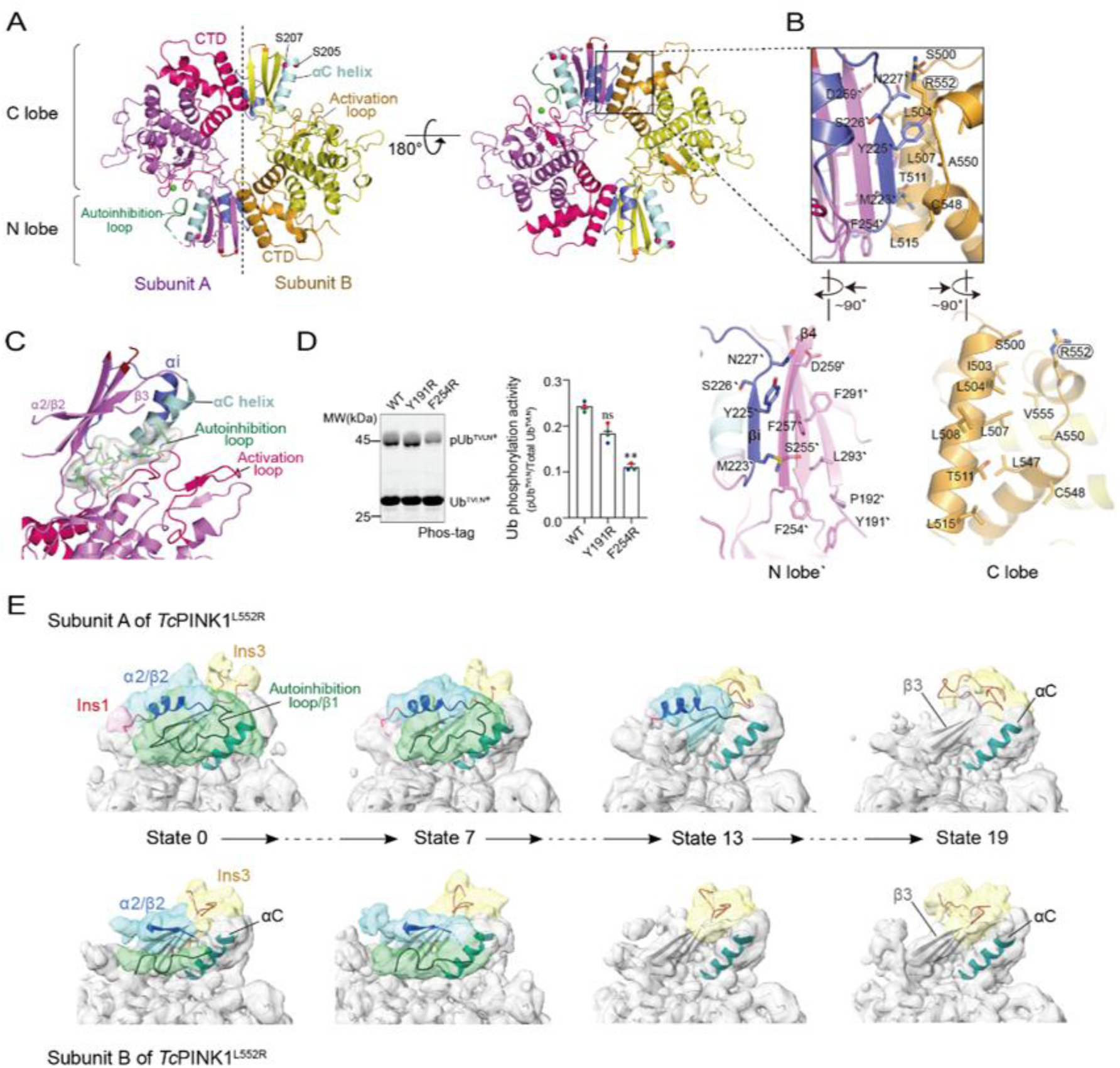
Two Distinct Conformations of the Dimeric Activating Mutant *Tc*PINK1^L552R^. **A)** Overall structure of dimeric *Tc*PINK1^L552R^ shown in cartoon representation. In this dimer, subunit A is auto-inhibited by the autoinhibition loop, while subunit B is priming to be activated. The αC helix is pale cyan; the autoinhibition loop is green; the αi helix is slate; activation loops are brown and hot pink; kinase domains are yellow and violet; CTDs are bright orange and hot pink. **B)** Interactions between the kinase N-lobe and CTD at the dimer interface. *Lower:* The interface is divided into two halves, with residues at the dimer interface shown as sticks. **C)** The autoinhibition loop is embedded in the active site of *Tc*PINK1^L552R^ in its autoinhibited state, displayed as green sticks and masked with a grey surface. **D)** Ub phosphorylation activities of WT and mutated *Tc*PINK1s assessed by fluorescent Phos-tag SDS-PAGE using the Ub^TVLN^ mutant, which is a more sensitive substrate for PINK1 phosphorylation (left). Statistics are based on triplicate technical repeats (right). **E)** 3D variability analysis of dynamic conformational changes in *Tc*PINK1^L552R^. Classified Cryo-EM density maps from 4 of 20 states are shown, contoured at 3σ. Refined low-resolution models are depicted in cartoon form. Segmented densities for Ins1, β1, β2/α2, and Ins3 are colored pink, green, cyan, and yellow, respectively, with the αC helix in deep teal.

In the autoinhibited state, two *Tc*PINK1 kinases form a symmetric, back-to-back dimer, where one kinase uses its C-terminal regulatory domain (CTD) to bind to the N-lobe of the other (Fig. 1A, 1B). These interactions stabilize both N-lobes of the two kinases in an active-like position relative to their C-lobes (Fig. S5, S6). The hydrophobic dimer interface buries a total surface area of 2712 Å². The interfacial residues at the CTD largely overlap with those forming the crystal contacts (Fig. S1), and the residues at the kinase N-lobe interface are highly conserved (Fig. S1). The surface of the dimer exhibits a bipolar electrostatic potential, with one side being more negatively charged and the other more positively charged (Fig. S5A).

However, the two subunits within this autoinhibited dimer adopt different conformations. In one subunit, the N-terminal fragment (residues 157-166) of the kinase, which includes part of the β1 strand, becomes partially disordered and embeds into the active site, blocking ATP and substrate access (Fig. 1C, Fig. S5B). The sequence of this autoinhibition loop is highly conserved, suggesting that a similar autoinhibitory mechanism might be adopted across species (Fig. S1). In the other subunit, the autoinhibition loop is released from the active site, accommodating ATP and substrate binding (Fig. 1A).

In the conformation primed for activation, the two kinases maintain the same dimeric interactions as in the autoinhibited state, but the autoinhibition loops are released from the active sites of both kinases (Fig. S5C). Additionally, the local conformation around the N-terminus of the αC helix is reorganized (Fig. S5D).

A detailed inspection of this primed activation conformation reveals several distinct features from early studies. First, the R-spine residues align similarly to the activated structure of Protein Kinase A (PKA) (Fig. S6), whereas the C-spine residues in the kinase N-lobe are aligned differently from early reported *Tc*PINK1 structures (Kumar, Tamjar et al. 2017, Okatsu, Sato et al. 2018). V176 moves away from the ATP binding site, while A194 on the β3 strand flips over, leaving the ATP binding site uncapped (Fig. S6A). Second, the kinase αC and αi helices are aligned parallel to each other and form extensive interactions along their entire lengths (Fig. S5D). Third, the β3 strand flips over, with K196 pointing toward the dimeric interface rather than the αC helix (Fig. S5C). The interactions between K196 and E217 on the αC helix were proposed previously to be essential for PINK1 activation (Kumar, Tamjar et al. 2017, Okatsu, Sato et al. 2018). However, compared to the K339A mutation, the K196A mutation only reduces but does not abolish the kinase activity of *Tc*PINK1^135-570^ (Fig. S5E).

3D variability analysis reveales the conformational dynamics of *Tc*PINK1^L552R^ during its transition between the autoinhibited and primed activation states (Fig. 1E). In the autoinhibited state, the autoinhibition loop is connected to the β2 strand, which can transform into an α helix under certain states (hereafter referred to as the α2/β2 helix/strand) (Mov. S1). During this transition, the autoinhibition loop disengages from the active site, leading to the dissociation of the α2/β2 helix/strand from the β3 strand. Accompanying these conformational changes, Insertion 3 becomes partially ordered. Additionally, in low-resolution structures refined with Rosetta, we found that Insertion 1 is located near the dimer interface (Fig. S5F) (Wang, Song et al. 2016).

To study the impact of this back-to-back dimerization on *Tc*PINK1^L552R^ activation, we mutated several conserved N-lobe residues at the dimer interface to charged residues to disrupt this dimerization (Fig. 1D; Fig. S5G, S5H). Except for the Y225R, Y225E and Y191R mutations, all other mutations abolish or diminish the ubiquitin phosphorylation activity of *Tc*PINK1. These results suggest that this back-to-back dimerization is potentially involved in regulating PINK1 activity.

Furthermore, we introduced the same mutations into *Tc*PINK1^L547R^, which is predominantly monomeric at low concentrations but can dimerize in a concentration-dependent manner (Fig. S5I). The Y191R mutation reduces the catalytic activity of the L547R mutant without affecting the wild-type (WT) *Tc*PINK1, while other mutations exhibit comparable effects on both the WT kinase and the L547R mutant (Fig. 1D, S5J). These data suggest that the N-lobe residues at the dimer interface do not only influence the dimeric assembly of PINK1 but are also crucial for its catalytic activity.

### The Phosphomimetic Mutant *Tc*PINK1^DDEE^ Forms an Inactive, Transitional Dimer

To understand the conformational transition of *Tc*PINK1, we determined the Cryo-EM structure of the phosphomimetic mutant *Tc*PINK1^DDEE^ (residues 150-570), which contains the S205D, S377D, T386E, and T530E mutations, at a resolution of 2.5 Å (Fig. S7, Table S1) (Okatsu, Sato et al. 2018). This mutant adopts a distinct conformation and dimeric interaction from *Tc*PINK1^L552R^.

Overall, *Tc*PINK1^DDEE^ forms a similar back-to-back dimer as *Tc*PINK1^L552R^ (Fig. 2A), with one subunit in the dimer rotating 40.4° and translating 12.3 Å relative to the other (Fig. S8A). The dimer surface is electrostatically bipolarized, similar to that observed in *Tc*PINK1^L552R^ (Fig. S8B). The residues contributing to the dimerization of *Tc*PINK1^DDEE^ largely overlap with those involved in the dimerization of *Tc*PINK1^L552R^ (Fig. 1B, 2B, S1), except for the β4 strand, which is flipped over and forms different interactions with the other subunit (Fig. 2B).

**Figure 2.**
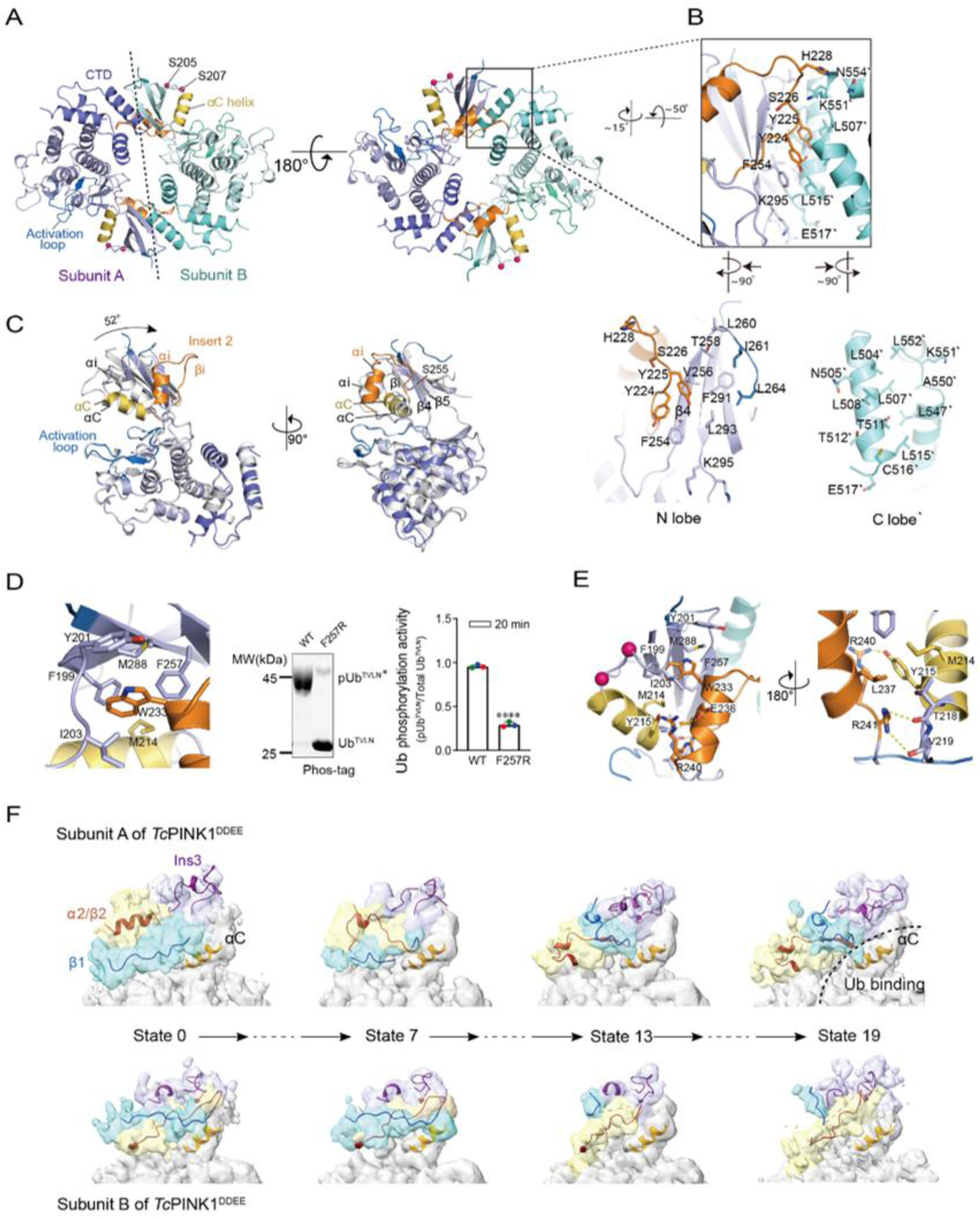
Dimeric Structure of the Phosphomimetic Mutant *Tc*PINK1^DDEE^. **A)** Overall structure of the dimeric phosphomimetic mutant *Tc*PINK1^DDEE^ shown in cartoon representation. The αC helix is yellow; the αi helix is bright orange; activation loops are blue and teal; kinase domains are pale blue and pale cyan; CTDs are slate and cyan green. **B)** Interactions between the kinase N- lobe and CTD at the dimer interface. *Lower:* The interface is divided into two halves, with residues at the dimer interface depicted as sticks. **C)** Interactions between the αi helix, αC helix, and N-lobe β strands. **D)** The F257R mutation stabilizes the structure of *Tc*PINK1^DDEE^ and reduces Ub phosphorylation activity, as determined by fluorescent Phos-tag SDS-PAGE using the Ub^TVLN^ mutant (left). Statistics are based on triplicate technical repeats (right). **E)** Interactions between the αC and αi helices at their C-termini in the *Tc*PINK1^DDEE^ structure. Hydrogen bonding interactions between the αC and αi helices are indicated with yellow dashes. Residues L237, Y215, and M214, shown in sticks, form hydrophobic interactions. **F)** 3D variability analysis of dynamic conformational changes in *Tc*PINK1^DDEE^. Classified Cryo-EM density maps from 4 of 20 states are shown, contoured at 3σ. Refined low-resolution models are depicted in cartoon form. Segmented densities for β1, β2/α2, and Ins3 are colored cyan, yellow, and violet-blue, respectively, with the αC helix in orange.

In addition to the dimerization interface, significant conformational changes are observed in each subunit at the kinase N-lobe and the Insertion 2 regions of *Tc*PINK1^DDEE^ (Fig. 2C). Compared to *Tc*PINK1^L552R^, the αi helix of *Tc*PINK1^DDEE^ is rotated 52° around its C-terminus and lifts along with the disordered βi strand. This lifting action pushes the partner kinase backward, allowing the tilting of the kinase N-lobe β-sheet and the swinging of the β4/β5 strands (Fig. 2A-C, S8A). In company with this series of conformational changes, the β4 strand flips over from S255 (Fig. 2B).

The elevated αi helix makes different interactions with the αC helix and the N-lobe β strands. At the N-terminus of the αi helix, W233 is buried within a conserved hydrophobic cavity (Fig. 2D), while at the C-terminus, R240 and R241 form hydrogen bonds with Y215, T218, and V219 (Fig. 2E).

Further 3D variability analysis reveals that *Tc*PINK1^DDEE^ also undergoes dynamic conformational changes (Fig. 2F, Mov. S2). During these transitions, the α2 helix unwinds and is transformed into the β2 strand, which subsequently aligns with the β3 strand. Meanwhile, the β1 strand tilts upward toward the dimer interface. In concert with these movements, Insertion 3 becomes disordered and then reorganizes with the β1 strand, creating a docking site for substrate recognition.

To assess the functional relevance of these structural changes, we measured the kinase activity of *Tc*PINK1^DDEE^. Consistent with previous reports, this mutant exhibit impaired kinase activity (Fig. S8C) (Okatsu, Sato et al. 2018). We then introduced an F257R mutation on the β4 strand to stabilize the conformation of *Tc*PINK1^DDEE^, as R257 could form π-cation interactions with residues W233, F199, and Y201 in this conformation (Fig. 2D). But this F257R mutation will disrupt the interface of *Tc*PINK1^L552R^-like dimer. Consistent with our structural prediction, the kinase activity of the F257R mutant is reduced relative to the WT *Tc*PINK1^135-570^ (Fig. 2D), suggesting that *Tc*PINK1^DDEE^ adopts an inactive, transitional conformation.

### Oligomeric Structures of *Tc*PINK1^DDEE^ Reveal Dimer-Dimer Interactions For PINK1 Trans-Autophosphorylation

In addition to monomers and dimers, *Tc*PINK1^DDEE^ also assembles into tetramers and hexamers in solution, as detected by Cryo-EM classification and analytical ultracentrifugation (AUC) (Fig. 3A- C). These higher-order oligomers are assembled as a chain of dimeric rings and exhibit similar dimer-dimer interactions (Fig. 3A, 3C). Analysis of these interactions of *Tc*PINK1^DDEE^ reveals a mechanism for trans-autophosphorylation.

**Figure 3.**
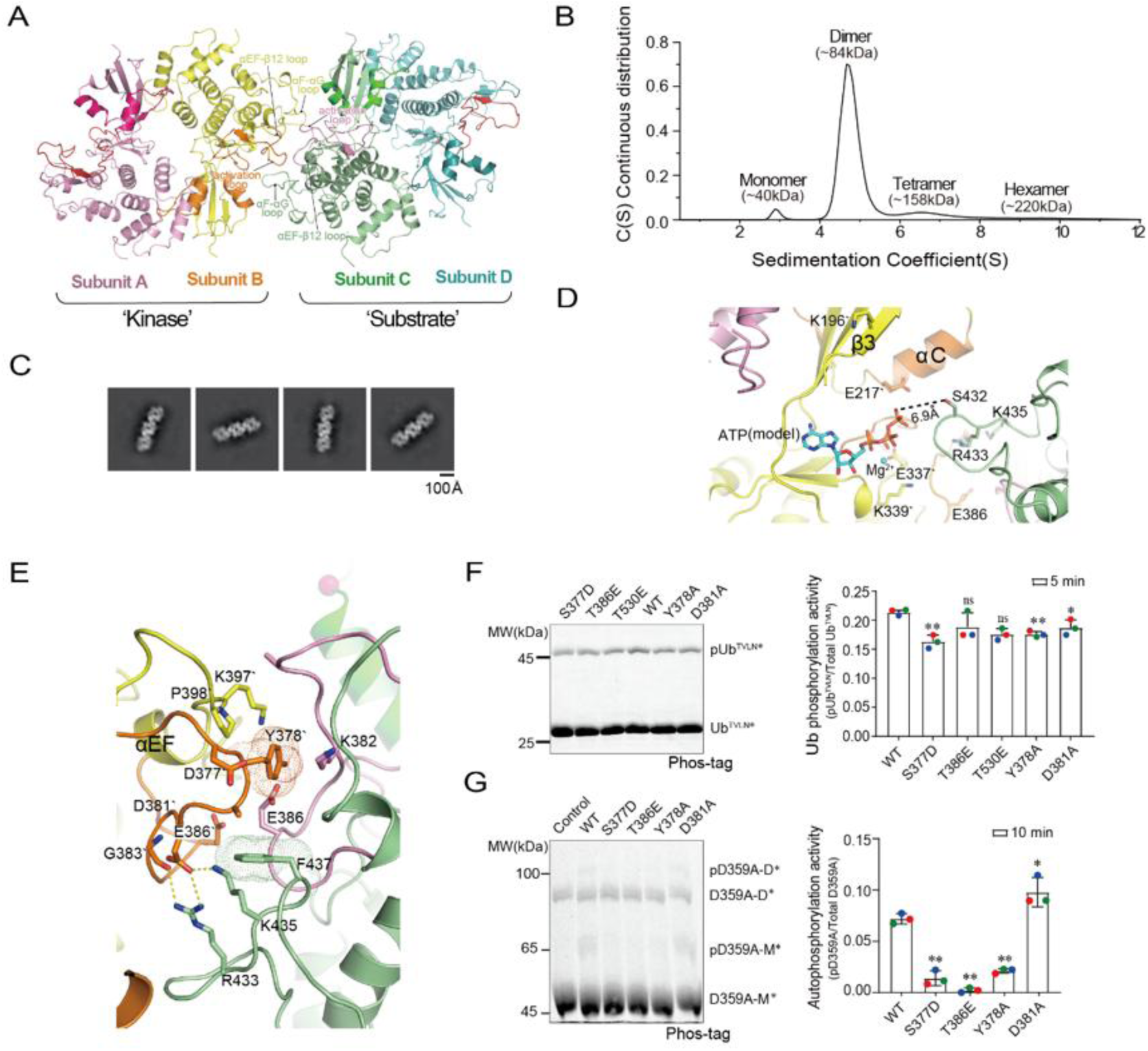
Structures of the Dimer-Dimer Trans-Autophosphorylation of *Tc*PINK1^DDEE^. **A)** Overall structure of the tetrameric trans-autophosphorylation complex of *Tc*PINK1^DDEE^. Based on ATP docking models, the two *Tc*PINK1^DDEE^ dimers are asymmetric, with one functioning as the ‘kinase dimer’ (subunits A and B) and the other as the ‘substrate dimer’ (subunits C and D). **B)** AUC analysis of purified *Tc*PINK1^DDEE^. In the c(S) distribution, four peaks corresponding to the molecular masses of 40, 84, 158, and 220 kDa were identified as monomers, dimers, tetramers, and hexamers of *Tc*PINK1^DDEE^ in solution. **C)** Representative 2D class averages of hexameric *Tc*PINK1^DDEE^. **D)** Modeled ATP binding mode during trans-autophosphorylation. ATP was modeled into the active site of subunit B by superimposing the structure of ANP-bound *Tc*PINK1 (PDB 5YJ9) onto this subunit. **E)** Interactions between the ‘kinase dimer’ and ‘substrate dimer’ at the trans-autophosphorylation interface. Residues involved in trans-autophosphorylation are shown as sticks, with two aromatic residues at the interface indicated by dots. **F)** Mutations at the dimer-dimer trans-autophosphorylation interface had a mild effect on the phosphorylation of Ub^TVLN^ by *Tc*PINK1, as shown by fluorescent Phos-tag SDS-PAGE (left) and statistics from triplicate technical repeats (right). **G)** Mutations at the dimer-dimer trans-autophosphorylation interface significantly altered the trans-autophosphorylation activity of *Tc*PINK1 toward the kinase-deficient mutant D359A, with results from fluorescent Phos-tag SDS-PAGE (left) and statistics from triplicate technical repeats (right).

The tetrameric structure of *Tc*PINK1^DDEE^ was resolved at a resolution of 3.52 Å (Table S1), revealing two back-to-back dimers in contact. Although some anisotropy in the density map was observed due to the preferred orientations of the tetrameric particles, the map was sufficiently clear to allow for atomic model building using the high-resolution structure of the dimeric *Tc*PINK1^DDEE^ as a reference.

In the tetramer, each dimer maintains a nearly identical structure to the dimeric *Tc*PINK1^DDEE^, with root mean square deviations (RMSDs) of Cα atoms measuring 0.67 Å and 0.73 Å, respectively. These dimers interact through their activation loops, specifically the αEF-β12 loop and the αF-αG loop (Fig. 3A). Through these dimer-dimer interactions, S432 from one subunit (either subunit B or C) is positioned in the active site of the other subunit, serving as a potential acceptor for trans- phosphorylation (Fig. 3D).

Due to the asymmetric nature of these dimer-dimer interactions, both dimers cannot simultaneously adopt a catalytically competent state. This was evident when we modeled ATP into the active sites of the kinases (Fig. 3D). In subunit C, the modeled ATP sterically interfered with residues from subunit B, while in subunit B, the modeled ATP did not encounter such steric hindrance. Moreover, in subunit B, the γ-phosphate of the modeled ATP was positioned 6.9 Å away from the hydroxyl group of S432 (Fig. 3D), aligning perfectly with the requirements for dissociative catalysis (Wang and Cole 2014). Consequently, we designated the dimer containing subunit B as the “kinase dimer” and the other as the “substrate dimer” (Fig. 3A). These dimers interact through salt bridges (D381/R433 and D381/K435), anion-π interactions (E386/Y378), and van der Waals contacts (Fig. 3E). Most interfacial residues in the “kinase dimer” are conserved, while those in the “substrate dimer” are variable (Fig. S1).

To investigate the impact of this dimer-dimer interaction, we employed the fluorescently labeled kinase-deficient mutant *Tc*PINK1^D359A^ as a substrate in autophosphorylation assays (Fig. S2, S3). Kinetic analysis revealed that WT *Tc*PINK1 phosphorylated *Tc*PINK1^D359A^ more efficiently than it did Ub, indicating a preferential mechanism for autophosphorylation (Table S2).

Subsequently, we introduced mutations at the “kinase dimer” interface to disrupt the binding of the substrate kinase. These mutations have minimal or mild effects on the Ub trans-phosphorylation activity of *Tc*PINK1 (Fig. 3F) but significantly impact the activity of *Tc*PINK1 in autophosphorylating *Tc*PINK1^D359A^ (Fig. 3G). This evidence confirms that the dimer-dimer interaction is critical for *Tc*PINK1 autophosphorylation (Fig. S8D).

Additionally, we evaluated whether S432 is a major phosphorylation site for *Tc*PINK1 by using the fluorescently labeled D359A/S432A double mutant. Compared to D359A, the phosphorylation level of D359A/S432A does not change significantly, indicating that S432 is not a major phosphorylation site for *Tc*PINK1 (Fig. S8E). Since all major autophosphorylation sites in this phosphomimetic mutant, *Tc*PINK1^DDEE^, have been mutated to Asp or Glu, it is likely that *Tc*PINK1^DDEE^ needs to engage a different autophosphorylation contact from early reported structures in phosphorylating minor sites, such as S432.

### The αA-CTD Interactions Are Not Essential for PINK1-Mediated Parkin Recruitment

It has been proposed that αA-CTD interactions are essential for PINK1 importation under mitochondrial stress (Kakade, Ojha et al. 2022, Rasool, Veyron et al. 2022, Callegari, Kirk et al. 2025). We therefore determined the Cryo-EM structure of WT *Tc*PINK1^117-570^ at a resolution of 3.5 Å, which reveals a distinct dimeric assembly from previously reported structures as well as the back-to-back dimer of *Tc*PINK1^L552R^ (Fig. 4A, S9, S10, Table S1). In this dimer, two kinases interact with each other with their αA and αK helices (Fig. 4A). Importantly, each kinase within this dimer adopts a fully active conformation: the activation loop is open; the essential triad for binding to the α and β phosphates (K196, E217) is well-aligned at the active site (Fig. 4A, S10); and the αC helix is kinked, with its N-terminal region positioned parallel to the αi helix from Insertion 2 (Fig. 4A, S6). Additionally, the C-spine and R-spine residues are strategically positioned to facilitate ATP binding and N-lobe positioning (Kornev et al., 2006; Kornev et al., 2008; Roskoski, 2016). The interactions between the αA helix and the CTD pull the β1-αA linker over and wrap it around the hydrophobic concave surface on top of the kinase N-lobe β sheet. Consequently, the β1 and β2 strands, along with the connecting G loop, are stabilized for ATP binding (Fig. 4A).

**Figure 4.**
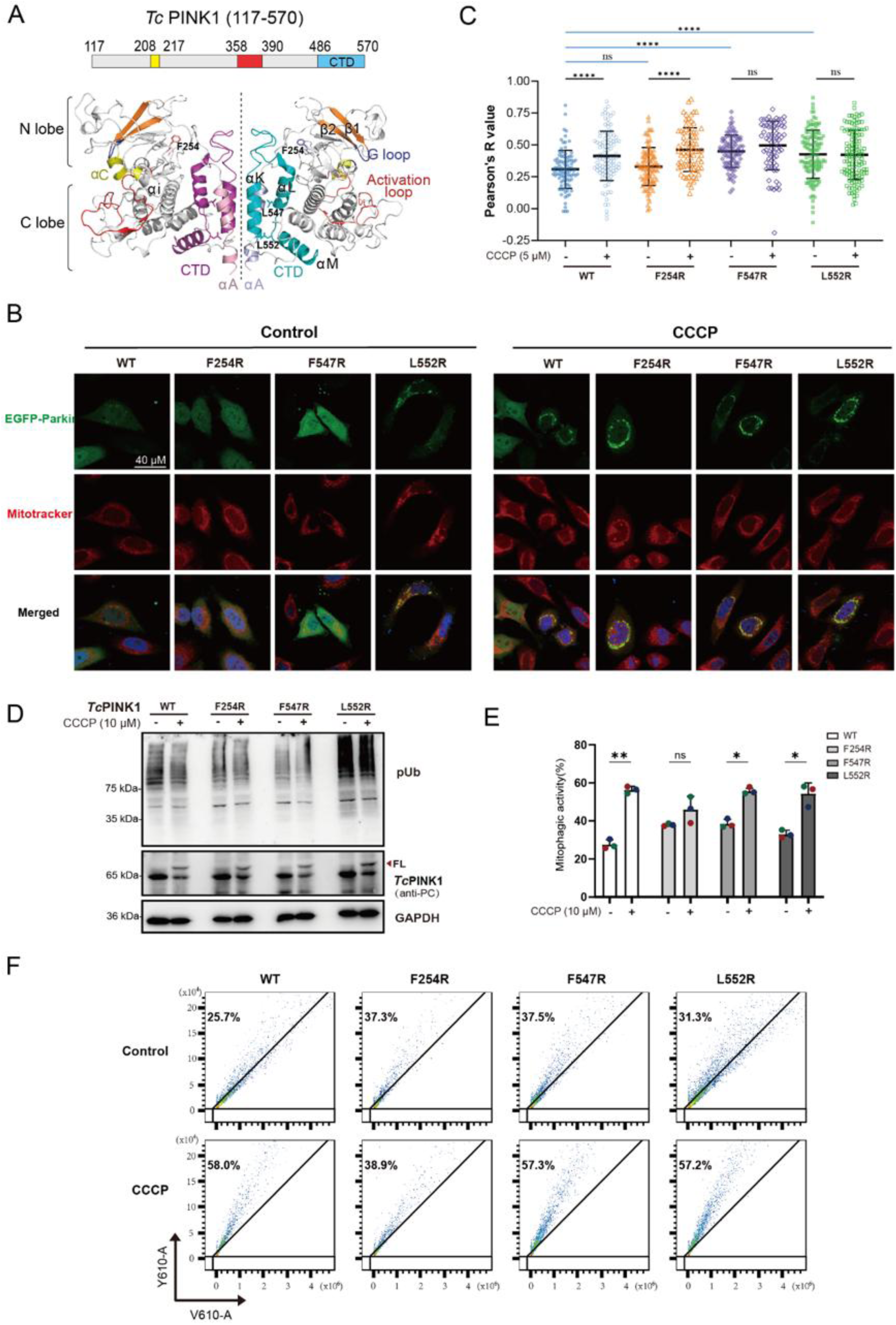
Effects of αA-CTD Interactions on Parkin Recruitment, Ub Transphosphorylation, and Mitophagy. **A)** *Tc*PINK1^117-570^ kinases form a symmetric dimer in solution through the interactions between their αA and αK helices. Top, diagram of the construct used in the Cryo-EM study. Bottom, the structure of *Tc*PINK1^117-570^ represented in cartoon. The αC helix, yellow; the activation loop, red; the CTDs, cyan and magenta; the αAs, light blue and pink. **B)** Parkin recruitment analyzed by confocal imaging. Shown are the representative confocal images of *h*PINK1 KO HeLa cells transiently transfected with plasmids encoding full-length *Tc*PINK1 WT or its mutants, along with EGFP-Parkin. Cells were treated with or without 5 µM CCCP for 1 hour prior to fixation. Mitochondria were stained with 100 nM Mitotracker, and nuclei were stained with 10 μg/mL DAPI. Images were captured using a Leica SP8 confocal microscope at 63× magnification. **C)** Statistical analysis of the colocalization between EGFP-Parkin and Mitotracker under various conditions. Over 70 images of EGFP-Parkin^+^ cells were randomly selected from replicated experiments for each sample and manually masked using ImageJ. The Pearson’s R value for colocalization of EGFP-Parkin and Mitotracker in each cell was calculated with the Coloc 2 program in ImageJ. Error bars represent the mean ± SD for each sample. Statistical significance is represented as: ns, no significance; *, p < 0.05; **, p < 0.01; ***, p < 0.001; ****, p < 0.0001. **D)** The expression, processing, and phosphorylation activities of WT *Tc*PINK1 and its mutants transfected transiently into *h*PINK1-KO COS7 cells. The transfected cells were treated with or without 10 μM CCCP for 8 hours and subsequently analyzed by Western blotting with antibodies against pUb, protein C, and GAPDH, respectively. The stabilized full-length *Tc*PINK1s are marked with a black arrow. **E,F)** The mitophagy activities of *Tc*PINK1 WT and mutants. Shown are the representative flowcytometry analysis of the mitophagy activities of *Tc*PINK1 transfected into *h*PINK1-KO HeLa cells treated with or without 10 μM CCCP for 10 hours (E). The statistics were determined from triplicated technical repeats, with error bars representing the mean ± SD (F).

However, in comparison to *Tc*PINK1^135-570^ (Fig. S3, Table S2), the overall turnover rate of *Tc*PINK1^117-570^ is slightly reduced (4.9 ± 0.3 *vs*. 7 ± 2 μM⁻¹·min⁻¹). Additionally, αA-CTD interactions shift the equilibrium of *Tc*PINK1^117-570^ toward monomers in solution, as shown by AUC analysis (Fig. S8F). These biochemical findings are incompatible with the notion that αA-CTD interactions stabilize *Tc*PINK1^117-570^ in an activated state, suggesting alternative regulatory mechanisms may be at play.

To investigate the impact of αA-CTD interactions on PINK1 importation and Parkin recruitment, we introduced three mutations (F254R, L547R, and L552R) into the full-length *Tc*PINK1 to modulate αA-CTD interactions. Based on the *Tc*PINK1^117-570^ structure, the F254R mutation is expected to stabilize the αA helix, while the L547R and L552R mutations are predicted to impair αA-CTD interactions (Fig. 4A). Subsequently, we transfected full-length WT *Tc*PINK1 and these mutants, along with EGFP-Parkin, into human (*h*)PINK1 knock-out (KO) HeLa cells, which do not express detectable levels of endogenous Parkin. In addition, the cellular systems and experimental conditions were pre-optimized and validated by using the WT *h*PINK1.

Comparing the results with WT *Tc*PINK1, the F254R mutation did not significantly affect CCCP (carbonyl cyanide m-chlorophenyl hydrazone)-induced Parkin recruitment, as evidenced by confocal imaging and Pearson’s co-localization R values (Fig. 4B, 4C). In contrast, in cells transfected with L552R and L547R, EGFP-Parkin was constitutively recruited to mitochondria, and CCCP treatment stimulated Parkin aggregation into puncta or mitochondria-derived vesicles (Fig. 4B, 4C).

In addition, we analyzed the expression, processing, and activities of transfected *Tc*PINK1s in *h*PINK1-KO COS7 cells. In the absence of CCCP, the expressed *Tc*PINK1s are detected as truncated products, whereas induction of mitochondrial depolarization leads to enhanced accumulation of the full-length *Tc*PINK1s, indicating *Tc*PINK1s are properly translocated into the mitochondria of these cells (Fig. 4D, S11A, S11B). Moreover, the expressed *Tc*PINK1s are active in Ub phosphorylation, with*Tc*PINK1^L552R^ mutant showing a trend of higher activity than the WT (Fig. 4D, S11A, S11B).

Consistently, in transfected *h*PINK1-KO HeLa cells, CCCP treatment could stimulate the mitophagy activities of WT *Tc*PINK1, the L552R, and L547R mutants but not the activity of the F254R mutant, confirming that the F254R mutant is less active in mitophagy (Fig. 4E, 4F, S11C). Collectively, these results suggest that the dimeric structure adopted by *Tc*PINK1^L552R^ is more favorable for Parkin recruitment.

## Discussion

To elucidate the molecular mechanisms governing PINK1 activation, we analyzed the structures and activities of *Tc*PINK1 in mitophagy. Utilizing our proposed criteria, we developed a sensitive fluorescent-based kinase activity assay, determined the Cryo-EM structures of *Tc*PINK1, and assessed its kinase and Parkin recruitment activities across various conditions and mutations. These results provide a comprehensive understanding of PINK1’s activation process in mitophagy.

In response to mitochondrial stress, multiple regulatory steps and distinctive functions of PINK1 are required and interlinked together to initiate mitophagy. Correspondingly, PINK1 is demanded to adopt different structural states to fulfill these functions, including the accumulation on the mitochondrial outer membrane, dimerization, auto-phosphorylation, Ub trans-phosphorylation, and Parkin recruitment (Lazarou, Jin et al. 2012, Okatsu, Uno et al. 2013). Early structural studies proposed that PINK1 forms a disulfide-linked face-to-face dimer on mitochondrial translocon in its priming to activation or autophosphorylation (Fig. S8G) (Gan, Callegari et al. 2022, Rasool, Veyron et al. 2022, Callegari, Kirk et al. 2025). However, this dimeric state is not competent for transphosphorylation of Ub and Parkin, which are prerequisite for the mitochondrial recruitment of Parkin and the initiation of mitophagy. Although it was proposed that the disulfide-linked dimer could be dissociated into active monomers under undefined conditions for mediating these phosphorylation events, at least for *Tc*PINK1, those monomers are equally incompetent for Parkin recruitment. Indeed, we identified *Tc*PINK1 as assembled into back-to-back dimers. Those dimers could not only trans-phosphorylate *Tc*PINK1 themselves, Ub, and Parkin, but are also able to mediate Parkin recruitment and mitophagy. However, our cellular studies are limited by using various overexpression systems. Further studies are therefore warranted to clarify the conformational status of PINK1 in its association with Parkin under physiologically relevant environment and conditions.

Early structural studies identified three conformations of PINK1: monomeric, ubiquitin-bound, and dimeric transphosphorylation states (Kumar, Tamjar et al. 2017, Schubert, Gladkova et al. 2017, Okatsu, Sato et al. 2018, Gan, Callegari et al. 2022, Rasool, Veyron et al. 2022, Callegari, Kirk et al. 2025). However, the complete transition from resting to stress-stimulated states remains elusive, leaving questions about how these conformations are related to PINK1’s diverse functionalities and overall activation landscape.

Using an active and a phosphomimetic mutant, we discovered that *Tc*PINK1 can adopt multiple dimeric conformations: autoinhibition, inactive transition, and priming to activation. In these states, *Tc*PINK1 forms a back-to-back dimer where its N-lobe interacts with its CTD. These dimers can trans-phosphorylate one another through dimer-dimer interactions and are crucial for PINK1-mediated Parkin recruitment under mitochondrial stress (Fig. 5). These findings suggest that: 1) phosphorylated *Tc*PINK1 forms a symmetric, back-to-back dimer essential for Ub phosphorylation and Parkin recruitment; 2) this dimer undergoes dynamic conformational changes; and 3) this dimer differs from the previously identified transphosphorylation dimer in activity and assembly. As suggested that the transphosphorylation dimer might be a priming state formed during PINK1 autophosphorylation, once phosphorylated, PINK1 may associate into a stable back-to-back dimer to mediate ubiquitin phosphorylation and Parkin recruitment under mitochondrial stress. Alternatively, these back-to-back dimers may assemble upon association with the TOM complex during mitochondrial importation, leading to PINK1 conformational changes, autophosphorylation via dimer-dimer interactions, Ub phosphorylation, and thereby Parkin recruitment (Fig. 5).

**Figure 5.**
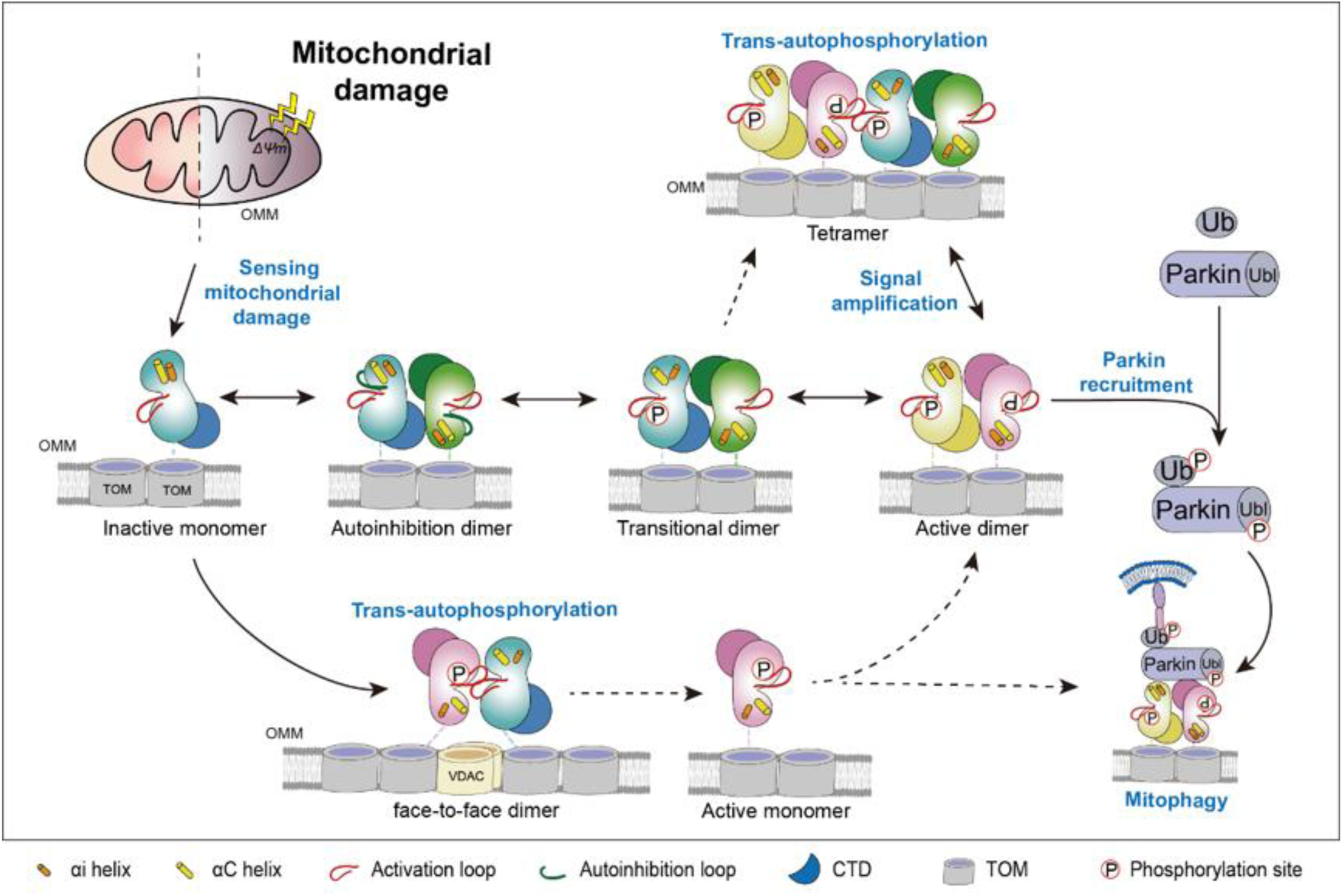
Model of Dynamic Conformations in the Activation of PINK1 Under Mitochondrial Stress. Under resting conditions, PINK1s are processed in mitochondria and degraded in cytosol. Upon sensing mitochondrial damage, PINK1 associates with the TOM complex and accumulates on the outer mitochondrial membrane. This accumulation allows monomeric, autoinhibited PINK1 molecules to auto-phosphorylate each other through transiently formed trans-phosphorylation dimers. Once phosphorylated, PINK1 organizes into stable back-to-back dimers, which mediate ubiquitin phosphorylation and Parkin recruitment under mitochondrial stress. Alternatively, these back-to-back dimers may form upon interaction with the TOM complex during stress, promoting conformational changes of PINK1, enabling its autophosphorylation through dimer-dimer interactions, and thereby enhancing Ub phosphorylation and Parkin recruitment. Within these back-to-back dimers, *Tc*PINK1 undergoes dynamic conformational changes across various states, including an autoinhibited state, an inactive transitional state, a primed activation state, and an active state. The autoinhibition loop, activation loop, αC helix, and αi helix are represented as green, red, yellow, and orange rods, respectively.

Supporting this possibility, existing evidence highlights the importance of TOM-associated PINK1 dimers for PINK1 activation and Parkin recruitment in mitophagy (Liu, Vives-Bauza et al. 2009, Lazarou, Jin et al. 2012, Okatsu, Uno et al. 2013). Using GFP- and mCherry-tagged PINK1s in the NAMOS assay, Okatsu *et al*. demonstrated that PINK1 dimerizes with the TOM complex under stress, correlating with its enhanced autophosphorylation activity and Parkin recruitment onto depolarized mitochondria (Okatsu, Uno et al. 2013). Mutations that impair PINK1 kinase activity or disturb its activation loop (S402A) deleteriously affect PINK1 importation and Parkin recruitment. In contrast, the S228A mutation on the αC helix, which mediates face-to-face dimerization, has minimal impact on either TOM association or Parkin recruitment. These evidence provide further support for our structural models.

As the αA helix of *h*PINK1 is critically involved in PINK1 mitochondrial importation(Kakade, Ojha et al. 2022, Rasool, Veyron et al. 2022, Callegari, Kirk et al. 2025), which is linked to its activation, we investigated the impact of αA-CTD interactions on PINK1-mediated mitophagy. In the structure of *h*PINK1-TOM complex, the αA helix interacts with PINK1 CTD, the TOM20, and the phosphate group of a bound lipid(Callegari, Kirk et al. 2025). As such, the αA helix restricts the movement of the kinase N-lobe and thus engages in the regulation of the kinase functions. However, the N-terminal fragment of αA helix is highly variable in length and amino acid composition (Fig. S1). This variability might indicate that the positioning of PINK1 on the TOM complex is flexible due to the plasticity of this juxtamembrane linkage. Consistently, Rasool *et al*. reported that the *Tc*PINK1 fragment, containing 121-160 a.a., is dynamically exposed to hydrogen-deuteron exchange in solution(Rasool, Soya et al. 2018). Alternatively, it is possible that the association between the N-terminal fragment of αA helix and CTD is not conserved so that αA could be displaced from the CTD to form a back-to-back dimer as seen in our study in its activation and/or Parkin recruitment. Moreover, αA-CTD interactions can be regulated under physiological conditions. During maturation, *h*PINK1 is cleaved by PARL at F104, yielding a 52 kDa product that is ubiquitinated at K137 (Liu, Guardia-Laguarta et al. 2017). Based on our *Tc*PINK1^117-570^ structure, ubiquitination at K137 could disrupt the interaction between the N-terminal αA helix and the CTD, indicating that αA-CTD interactions can be modulated under physiological conditions.

Through 3D variability analysis, we captured dynamic conformational transitions of both activating and phosphomimetic mutants. Notably, the activating mutant exhibited dual distinct conformations, suggesting complex regulatory potential for PINK1 assembly and kinase activity. This complexity mirrors the conformational dynamics observed in Abl kinase, which relate to drug resistance (Xie, Saleh et al. 2020). Our findings provide critical structural insights into PINK1’s conformational plasticity, prompting further investigation into its implications for Parkinson’s disease pathogenesis and treatment.

Mounting evidence suggested that lipids are critically involved in the quality control and clearance of mitochondria(Chu, Ji et al. 2013, Oleinik, Albayram et al. 2023, Feng, Cai et al. 2025). For example, cardiolipin, a negatively charged lipid, is translocated to the mitochondrial outer membrane upon the initiation of mitophagy, which accompanies the activation of PINK1 and the recruitment of Parkin to mitochondria(Chu, Ji et al. 2013, Shen, Li et al. 2017, Feng, Cai et al. 2025). Indeed, in the structure of *h*PINK1-TOM complex, the *h*PINK1 R119 at the membrane interface interacts with the phosphate group of a bound lipid(Callegari, Kirk et al. 2025). However, this positively charged N-terminal fragment is not conserved in *Tc*PINK1 as well as some other species (Fig. S1). Given the electrostatically polarized surface of *Tc*PINK1 back-to-back dimer, we tempt to speculate that *Tc*PINK1 dimer might be oriented and stabilized on the mitochondrial outer membrane with its positively charged convex side interacting closely with those negatively charged lipids (Fig. S5A, S8B, S8D). As such, multiple signals of mitochondrial damage can be concertedly integrated to regulate the activation of PINK1-Parkin axis in mitophagy. However, such a possibility needs to be extensively validated in future studies. In summary, our study lays the foundation for understanding the allosteric regulation of *Tc*PINK1 kinase activity through its dimerization.

## Materials and Methods

### Plasmid construction

The genes encoding *Tc*PINK1 (a.a. 117–570), *Tc*PINK1 (a.a. 135–570), and *Tc*PINK1^DDEE^ (a.a. 153–570, with S205D/S377D/T386E/T530E mutations), along with other mutants fused to an N-terminal His×6-tag, were subcloned into the pET-15b vector at the *Bam*HI and *Xho*I sites. The genes for His×6-TEV-GGG-SUMO-fused Ub and Ub^TVLN^ were inserted into the pET-28a vector at the *Nhe*I and *Xho*I sites. Full-length *Tc*PINK1 and its mutants (L552R, L547R, and F254R) were subcloned into the pEF-1 vector via homologous recombination, while the gene for EGFP-Parkin was subcloned into the pcDNA3.1 vector using the same method. The plasmid encoding the mt-Keima was generously provided by Dr. Yufeng Yang (Fuzhou University) as a gift.

### Purification of *Tc*PINK1 (117-570), *Tc*PINK1 (135-570) and *Tc*PINK1^DDEE^ (153-570)

*E. coli* BL21 (DE3) cells were transformed with pET-15b plasmids encoding *Tc*PINK1 (117-570), *Tc*PINK1 (135-570), and *Tc*PINK1^DDEE^ (153-570). The transformed cells were cultured in LB medium with 100 mg/L ampicillin at 37 °C until the optical density at 600 nm (OD_600_) reached 0.6- 0.8. Protein expression was induced by adding 0.4 mM IPTG, and the cells were cultured for an additional 18 h at 16 °C.

Cells were harvested by centrifugation at 4,000 rpm for 15 min and lysed using a French Press in lysis buffer (50 mM Tris-HCl (pH 8.0), 500 mM NaCl, 20 mM Imidazole, 10% Glycerol, 0.5% Triton X-100, and 1 mM PMSF). The lysate was clarified by centrifugation at 18,000 rpm for 30 min at 4 °C. The supernatant was loaded onto a Ni^2+^-NTA column, washed with binding buffer (50 mM Tris-HCl (pH 8.0), 500 mM NaCl, 20 mM Imidazole, 10% Glycerol), and eluted with elution buffer (50 mM Tris-HCl (pH 8.0), 500 mM NaCl, 300 mM Imidazole, 10% Glycerol).

Affinity-purified proteins were buffer-exchanged using a 5 mL Hitrap Desalting column (Cytiva) equilibrated in 20 mM Tris-HCl (pH 8.0), 20 mM NaCl, 5% Glycerol, and 1 mM DTT. Proteins were further purified by anion exchange chromatography on a 5 mL Hitrap Q column (Cytiva), equilibrated with 20 mM Tris-HCl (pH 8.0), 1 mM DTT, and 5% Glycerol, and eluted with a linear NaCl gradient (0 M to 1 M).

Subsequent size-exclusion chromatography was performed using a Superdex 200 Increase 10/300 GL column (Cytiva) equilibrated with 20 mM Tris-HCl (pH 8.0), 100 mM NaCl, and 1 mM DTT. Fractions containing the target proteins were pooled, concentrated, and supplemented with 20% (v/v) glycerol before being flash-frozen in liquid nitrogen and stored at -80 °C.

Other *Tc*PINK1 mutants were purified using the same protocol, with IPTG induction concentrations of 0.2 mM for *Tc*PINK1^Y191R^ and *Tc*PINK1^F193R^, and 0.1 mM for *Tc*PINK1^DDEE^. **Purification of Ubiquitin (WT Ub and Ub^TVLN^ mutant)** *E. coli* Rosetta (DE3) cells were transformed with plasmids encoding WT Ub or the Ub^TVLN^ mutant. The transformed cells were cultured in LB medium supplemented with 50 mg/L ampicillin and 34 mg/L chloramphenicol at 37 °C until the OD_600_ reached 0.6-0.8. Protein expression was induced by adding 1 mM IPTG, and the cells were incubated for an additional 5 h at 37 °C.

Cells were harvested by centrifugation at 4,000 rpm for 15 min and lysed via sonication in lysis buffer (20 mM Tris-HCl (pH 8.0), 400 mM NaCl, 20 mM Imidazole, 10% Glycerol, and 1 mM PMSF). The lysate was clarified by centrifugation at 18,000 rpm for 30 min at 4 °C. The supernatant was loaded onto a Ni^2+^-NTA column, washed with binding buffer (20 mM Tris-HCl (pH 8.0), 400 mM NaCl, 40 mM Imidazole, 10% Glycerol), and eluted with elution buffer (20 mM Tris-HCl (pH 8.0), 400 mM NaCl, 300 mM Imidazole, 10% Glycerol).

Affinity-purified proteins were buffer-exchanged using a 5 mL Hitrap Desalting column (Cytiva) equilibrated in 20 mM Tris-HCl (pH 8.0), 20 mM NaCl, 5% Glycerol, and 1 mM DTT. Further purification was performed using anion exchange chromatography on a 5 mL Hitrap Q column (Cytiva), equilibrated with 20 mM Tris-HCl (pH 8.0), 1 mM DTT, and 5% Glycerol, and eluted with a linear NaCl gradient from 0 M to 1 M.

The final purification step involved size-exclusion chromatography using a Superdex 200 Increase 10/300 GL column (Cytiva), equilibrated with 20 mM Tris-HCl (pH 8.0), 100 mM NaCl, and 1 mM DTT. Fractions containing the target proteins were pooled, concentrated, and supplemented with 20% (v/v) glycerol. The protein solution was then flash-frozen in liquid nitrogen and stored at -80 °C.

### *In vitro* ubiquitin phosphorylation assays

#### Phosphorylation of Ub

Purified and fluorescently-labeled Ub at a concentration of 14.88 μM was phosphorylated by 0.3 μM of either purified WT or mutant *Tc*PINK1 (135–570) at 30 °C in a reaction buffer containing 50 mM Tris-HCl (pH 7.5), 10 mM MgCl_2_, 2 mM ATP, 10 mM DTT, and 0.1% Triton X-100. After 10 or 20 minutes, the reaction was terminated by adding 6x SDS sample buffer.

Phosphorylated Ub was separated from non-phosphorylated Ub by electrophoresis on a 12% SDS-polyacrylamide gel, supplemented with 50 μM PhosBind Acrylamide (APExBIO) and 200 μM MnCl_2_. Fluorescent images of the Phos-tag SDS-PAGE gels were captured using a GE LAS-4000 imager, and the fluorescent intensity of the Ub bands was quantified using ImageJ. Data analysis was performed with Origin software.

#### Phosphorylation of Ub^TVLN^

For the phosphorylation of Ub^TVLN^, 0.03 μM WT or mutant *Tc*PINK1 (135–570) was pre-autophosphorylated in the reaction buffer at 30 °C for 10 minutes.

Subsequently, 30.0 μM of labeled Ub^TVLN^ was phosphorylated by the pre-autophosphorylated *Tc*PINK1 at 30 °C for 5 minutes. The experimental procedures and quantifications were conducted similarly to those in the Ub phosphorylation assay.

#### Analytical ultracentrifugation (AUC)

*Tc*PINK1 and its mutants were purified following previously established protocols. To compare the sedimentation rates of different mutants, protein samples were prepared at a concentration of 11.1 μM and subjected to centrifugation at 42,000 rpm using a Beckman centrifuge for 8 hours. After centrifugation, ultraviolet absorption data and interferometric optical scanning data were collected. The data were subsequently processed using Setfit software (Schuck 2000) to analyze sedimentation characteristics.

#### Labeling of Ubiquitin with FITC

Labeling was performed at 4 °C in a 500 μL reaction mixture containing 20 mM Tris-HCl (pH 8.0), 100 mM NaCl, 50 mM CaCl_2_, along with FITC-LPETGG, Ub, and Sortase A in a molar ratio of 250:50:1. The reaction was allowed to proceed for 16 hours.

After labeling, the protein sample was cleared by centrifugation at 12,000 rpm for 10 min at 4 °C. Labeled proteins were purified using a Hitrap Desalting column (Cytiva) equilibrated with 20 mM Tris-HCl (pH 8.0), 20 mM NaCl, and 5% Glycerol. Gel-filtration chromatography indicated that both unlabeled and labeled Ub were eluted at the same position with similar profiles.

To assess labeling efficiency, calibration curves for FITC and protein concentrations were generated. FITC concentration was determined by measuring the fluorescent intensity at an excitation wavelength of 495 nm and an emission wavelength of 515 nm as a function of FITC-LPETGG concentration. Protein concentrations were measured using the Bradford assay with BSA as the standard. The same labeling protocol was applied to Ub^TVLN^ and *Tc*PINK1^D359A^ (135- 570).

### Determination of reaction kinetics for Ubiquitin phosphorylation by *Tc*PINK1 (117-570), *Tc*PINK1 (135-570), and their mutants

#### Phosphorylation of Ub

Purified and FITC-labeled ubiquitin was serially diluted 2-fold in a dilution buffer containing 20 mM Tris-HCl (pH 8.0) and 100 mM NaCl. The diluted ubiquitin at final concentrations ranging from 74.3 μM to 0.04 μM was incubated with and phosphorylated by 0.3 μM WT *Tc*PINK1 (135–570) or its mutants at 30 °C in a reaction buffer containing 50 mM Tris-HCl (pH 7.5), 10 mM MgCl_2_, 2 mM ATP, 10 mM DTT, and 0.1% Triton X-100. After 5 or 10 minutes, the phosphorylation reaction was terminated by adding 6x SDS sample buffer.

Phosphorylated ubiquitin was separated from non-phosphorylated ubiquitin using 12% SDS-polyacrylamide gels containing 50 μM PhosBind Acrylamide (APExBIO) and 200 μM MnCl_2_. The Phos-tag SDS-PAGE gels were imaged with a GE LAS-4000 imager, and the fluorescent bands of ubiquitin were quantified using ImageJ. The data were processed with Origin software and fitted to the Michaelis-Menten equation. Means and standard deviations for the kinetic parameters were calculated from triplicate technical repeats.

#### Phosphorylation of Ub^TVLN^

For the phosphorylation of Ub^TVLN^, 0.01 μM WT *Tc*PINK1 (135-570) was pre-autophosphorylated in the reaction buffer at 30 °C for 10 minutes. Labeled Ub^TVLN^ at concentrations ranging from 60.4 μM to 0.03 μM was then phosphorylated by the pre- autophosphorylated WT *Tc*PINK1 (135-570) at 30 °C for 5 minutes. The experimental procedures were conducted similarly to those for ubiquitin phosphorylation.

To assess the reaction rate of Ub^TVLN^ phosphorylation by *Tc*PINK1 (117-570), 0.01 μM *Tc*PINK1 (117-570) was used to phosphorylate 10.0 μM FITC-labeled Ub^TVLN^. Reactions were stopped at specified time points, and the phosphorylated samples were analyzed as described. The reaction rate remains linear for up to 20 minutes. Subsequently, the concentrations of FITC-labeled Ub^TVLN^ were varied from 60.8 μM to 0.03 μM, and these samples were phosphorylated by 0.01 μM WT *Tc*PINK1 (117-570). After 5 minutes, the samples were separated by Phos-tag SDS-PAGE, and the kinetics of the reaction were analyzed as previously described.

#### *In vitro Tc*PINK1 autophosphorylation assays

Autophosphorylation assays were conducted at 30 °C in a 500 μL reaction mixture containing the appropriate buffer and substrates. For the pre- autophosphorylation step, 0.03 μM of either WT *Tc*PINK1 or its mutants were incubated for 10 minutes.

Following pre-autophosphorylation, the trans-autophosphorylation activity was assessed by adding approximately 51 μM *Tc*PINK1^D359A^ (135-570) to the reaction mixture and incubating for an additional 10 minutes. This step allowed for the comparison of trans-autophosphorylation activities between WT *Tc*PINK1 and its mutants.

To evaluate kinetic parameters, *Tc*PINK1^D359A^ was serially diluted to achieve concentrations ranging from 52.75 μM to 0.103 μM. Each diluted sample was phosphorylated using the pre- autophosphorylated 0.03 μM WT *Tc*PINK1 for 10 minutes at 30 °C. The experiments and quantifications followed the same methodology as the ubiquitin phosphorylation assays.

For the labeling and phosphorylation of *Tc*PINK1^D359A/S432A^, the same procedures employed for *Tc*PINK1^D359A^ were applied.

### Cryo-EM sample preparation and data acquisition

For Cryo-EM sample preparation, 4 μL of *Tc*PINK1(117-570) (0.45 mg/mL), *Tc*PINK1^L552R^ (0.25 mg/mL), and *Tc*PINK1^DDEE^ (0.4 mg/mL) were applied to holy Ni-Ti grids (R1.2/1.3, 300 mesh, Au, X-Pivot, China) glow-discharged with a Gatan Solarus Plasma Cleaning System 955. The grids were blotted for 1 second and flash-frozen in liquid ethane cooled by liquid nitrogen with a Vitrobot Mark IV (Thermo Fisher Scientific Inc.) set to 4 °C and 100% humidity.

Following freezing, the grids were transferred to a Titan Krios electron microscope (Thermo Fisher Scientific Inc.) operating at 300 kV. The microscope was equipped with a Gatan K3 Summit direct electron detector and a GIF Quantum energy filter. Movie stacks were collected automatically using EPU, with a preset defocus range from -1.2 μm to -1.8 μm in super-resolution mode. Data collection was conducted at a normal magnification of 130,000, resulting in a pixel size of 0.668 Å/pixel on the specimen. Each movie stack consisted of 32 frames and was exposed for 1.11 seconds. The slit width on the energy filter was set to 20 eV, with a total dose of approximately 50 e^-^/Å^2^ for each micrograph stack.

#### Image processing and analysis

Image processing was conducted using cryoSPARC (Punjani, Rubinstein et al. 2017), with strategies outlined in Fig. S4, S7, and S9. The movie stacks underwent motion correction via MotionCor2 (Zheng, Palovcak et al. 2017), and defocus values were estimated using Patch CTF estimation. Micrographs contaminated or with resolutions below 5 Å were excluded, resulting in 18,161 micrographs for *Tc*PINK1(117-570), 8,111 micrographs for *Tc*PINK1^L552R^, and 6,054 micrographs for *Tc*PINK1^DDEE^ used for structure determination.

#### *Tc*PINK1 (117-570) Dataset

For the *Tc*PINK1 (117-570) dataset, particles were auto-picked using blob picker and extracted from 500 micrographs with a box size of 400 pixels. These particles were classified into 50 classes through 2D classification. A total of 19,832 particles with representative 2D class averages were selected as the training dataset for Topaz Train (Bepler, Morin et al. 2019). Subsequently, 4,995,315 particles were auto-picked from the 18,161 micrographs using Topaz. After two rounds of 2D classification, 3,693,128 particles were selected and subjected to *Ab-initio* Reconstruction into five classes, which served as templates for Heterogeneous Refinement of all selected particles. Post-refinement, particles with detailed features and the highest resolution were chosen for a second round of *Ab-initio* Reconstruction and Heterogeneous Refinement. Non-uniform Refinement improved the density resolution to 4.07 Å (Punjani, Zhang et al. 2020, Zivanov, Nakane et al. 2020). Following C2 symmetry application and Local CTF Refinement, a final map with a resolution of 3.48 Å was obtained. All reported resolutions were determined using the gold-standard FSC 0.143 criterion, with local-resolution distributions evaluated using ResMap (Kucukelbir, Sigworth et al. 2014).

#### *Tc*PINK1^L552R^ Dataset

For the *Tc*PINK1^L552R^ dataset, a total of 80,034 particles were auto-picked using blob picker and extracted from 500 micrographs with a box size of 400 pixels. These particles were classified into 50 classes through 2D classification. A training dataset for Topaz Train comprised 1,827 particles from 100 micrographs with representative 2D class averages.

Using Topaz, 1,861,310 particles were auto-picked from 8,111 micrographs. After one round of 2D classification, 1,724,632 particles were selected and randomly split into two sets for *Ab-initio* Reconstruction in four classes, serving as templates for Heterogeneous Refinement. After refinement, 658,543 particles were selected from density maps with detailed features and the highest resolution. A second round of 2D classification refined this to 554,347 particles. These were further refined using CTF refinement and Non-uniform Refinement, achieving a density resolution of 3.02 Å (Punjani, Zhang et al. 2020, Zivanov, Nakane et al. 2020). A 3D variability analysis was performed to separate individual conformations, resulting in maps with 3.04 Å resolution for clusters representing the ‘prime-activation’ conformation, and 3.22 Å and 3.35 Å resolutions for clusters representing the ‘autoinhibited’ conformation.

#### *Tc*PINK1^DDEE^ Dataset

For the *Tc*PINK1^DDEE^ dataset, 151,416 particles were auto-picked using blob picker and extracted from 500 micrographs with a box size of 480 pixels. A training dataset for Topaz Train consisted of 7,189 particles from 100 micrographs with representative 2D class averages. Using Topaz, 1,570,982 particles were auto-picked from 6,054 micrographs. Following one round of 2D classification, 1,555,299 particles were subjected to *Ab-initio* Reconstruction in four classes, which were used as 3D volume templates for Heterogeneous Refinement. Post-refinement, two density maps with detailed features for the dimer and tetramer (30.5% for dimer and 27.0% for tetramer) were selected. The dimer class underwent a second round of *Ab-initio* Reconstruction, generating four models that were combined with the dimer map from the previous Heterogeneous Refinement for further refinement. This process yielded a map with a resolution of 2.77 Å (Bepler, Morin et al. 2019). Following Local CTF Refinement and Non-uniform Refinement, a final map with a resolution of 2.63 Å was obtained (Punjani, Zhang et al. 2020, Zivanov, Nakane et al. 2020). A 3D variability analysis in simple mode produced 20 low-resolution density maps for movie creation. After applying C2 symmetry, a cryo-EM map at 2.51 Å resolution was achieved. For the *Tc*PINK1^DDEE^ tetrameric class, particles were selected from 100 micrographs as a training dataset for Topaz. A total of 2,409,694 particles were auto-picked from 6,054 micrographs. These particles underwent *Ab-initio* Reconstruction and two rounds of Heterogeneous Refinement using the previous four volume maps as templates, resulting in a final density map with a resolution of 3.47 Å (Punjani, Zhang et al. 2020, Zivanov, Nakane et al. 2020).

### Model building and refinement

To generate initial models, the structures of *Tc*PINK1 kinase (PDB ID 5YJ9 or 7MP8) were docked into the Cryo-EM density maps using Chimera (Pettersen, Goddard et al. 2004). The models were iteratively rebuilt in COOT and refined using Phenix (Emsley, Lohkamp et al. 2010, Liebschner, Afonine et al. 2019). The geometry of the refined models was validated with MolProbity (Chen et al., 2010), while the registers were verified using Q-scores and CheckMySequence (Pintilie, Zhang et al. 2020, Chojnowski 2022). Statistics for the Cryo-EM data collection and model refinement are reported in Table S1. The refined coordinates and cryo-EM data have been deposited in the PDB and EMDB, respectively.

Additionally, the low-resolution Cryo-EM density maps generated from 3D variability analysis were utilized in the refinement of structural models with Rosetta, as previously described (Wang, Song et al. 2016, Punjani and Fleet 2021).

### Using confocal microscopy to characterize the recruitment of Parkin to mitochondria

For the analysis of αA-CTD interactions, *h*PINK1 knockout (*h*PINK1-KO) HeLa cells were cultured in DMEM supplemented with 10% FBS in 35 mm confocal dishes at 37 °C in a CO_2_ incubator.

Prior to transfection, the culture media was replaced with fresh media. Cells at approximately 70% confluency were transiently transfected with 2.5 μg of plasmid encoding full-length *Tc*PINK1 or its mutants, along with 2.5 μg of EGFP-Parkin plasmid, using 15 μg of PEI per well. After 4 hours, the culture media was replaced with fresh media. Two days post-transfection, cells were treated with or without 5 µM CCCP for 1 hour before fixation. Mitochondria were stained with 100 nM MitoTracker Deep Red FM (Yeasen Biotechnology), and nuclei were stained with 10 μg/mL DAPI, following the manufacturer’s instructions.

Images were captured using a Leica SP8 confocal microscope with a HC PL APO CS2 63x/1.40 OIL objective at 23 °C. Three-channel images were sequentially collected using 405 nm, 488 nm, and 638 nm lasers to detect signals from DAPI, EGFP-Parkin, and MitoTracker, respectively.

To assess the recruitment of Parkin to mitochondria, Pearson’s R values were calculated to quantify the co-localization of EGFP-Parkin and MitoTracker. The boundary of each EGFP-Parkin-positive cell (green channel) was individually masked. The Pearson’s R value for co- localization between EGFP-Parkin (green channel) and MitoTracker (red channel) in each cell was determined using the Coloc 2 program in ImageJ software (NIH). Confocal images were collected from triplicated transfection experiments, and the distribution and statistics of the Pearson’s R values were quantified from over 70 randomly selected cells for each sample.

### The flowcytometry analysis of the mitophagy activities of WT *Tc*PINK1 and its mutants

Equal amount of the plasmid encoding mt-Keima and the plasmid encoding one of the *Tc*PINK1^WT^, *Tc*PINK1^L552R^, *Tc*PINK1^F254R^, and *Tc*PINK1^L547R^ were transfected respectively into *hPINK1*-KO Hela cells with Lipo8000. The transfected cells were incubated at 37℃ in a 5% CO_2_ incubator for 24 hours. The culture media was replaced with fresh media. Then, the cells were treated with or without 10 µM CCCP for 10 hours. After treatment, the cells were washed once with PBS, detached with trypsin, and resuspended in PBS for FACS analysis. The FACS data were recorded with Beckman Coulter CytoFLEX and analyzed with FlowJo. The forward and side scattering of the cells were used in the gating of live cells. The mitophagy activities of the cells were determined by using a dual-color fluorescent distribution: one was set at 405 nm excitation and 610 nm emission, and the other was set at 561 nm excitation and 610 nm emission. The mock-transfected cells were used as a negative control in the gating of the cells. The population of the cells undergoing mitophagy was gated as described(Lin, Chen et al. 2020). The statistics were calculated from the results of triplicated biological repeats.

### Western blotting analysis of the expression and Ub phosphorylation activities of *Tc*PINK1s

*hPINK1*-KO COS7 cells were seeded into 12-well plates and cultured in DMEM supplemented with 10% FBS. Upon reaching 50∼60% confluency, the cells were transfected respectively with the plasmids encoding *Tc*PINK1^WT^, *Tc*PINK1^L552R^, *Tc*PINK1^F254R^, and *Tc*PINK1^L547R^ using Lipo8000. The transfected cells were cultured 24 hours before the culture media were replaced with fresh media. Then, the cells were treated with or without 10 µM CCCP for 8 hours. Next, the cells were washed twice with PBS and lysed with 100 µL/well lysis buffer (20 mM Tris-HCl, pH 8.0, 150 mM NaCl, 1 mM EDTA, 1 mM EGTA, 1% NP-40, 1% Sodium deoxycholate, 2.5 mM Sodium pyrophosphate, 1 mM beta-glycerophosphate, 1 mM Na_3_VO_4_, 1 mM PMSF, and 1x cocktail protease inhibitors). The lysates were cleared by centrifugation at 12,000 rpm for 10 minutes at 4 ℃. The supernatants were transferred into 1.5 mL Eppendorf tubes and mixed with 5x SDS loading buffer. These samples were separated by reducing SDS-PAGE followed by Western blotting with antibodies against pUb, protein C, and GAPDH (pUb, CST, cat # 62802S; protein C, GeneScript, cat # A00637; GAPDH, proteintech, cat # 60004-1-Ig). The intensities of the Western blotting bands were quantified with ImageJ. The statistics were determined from the results of triplicated biological repeats.

### Quantification and statistical analysis

#### Transphosphorylation Assays

For transphosphorylation assays, phosphorylated Ub, Ub^TVLN^, *Tc*PINK1^D359A^, or *Tc*PINK1^D359A/S432A^ were separated from their non-phosphorylated counterparts using 6-12% SDS-polyacrylamide gel, supplemented with 50 μM PhosBind Acrylamide (APExBIO) and 200 μM MnCl_2_. Fluorescent images of the Phos-tag SDS-PAGE gels were captured using a GE LAS-4000 imager, and the fluorescent bands were quantified with ImageJ. Means and standard deviations were calculated from triplicate experiments, and statistical significance was assessed using a paired T-test (ns: no significance; *, p < 0.05; **, p < 0.01; ***, p < 0.001; ****, p < 0.0001). Data were processed and presented using Origin.

#### Kinetic Experiments

In kinetic experiments, fluorescent images of the Phos-tag SDS-PAGE gels were captured with a GE LAS-4000 imager, and the fluorescent bands were quantified with ImageJ. The data were processed using Origin and fitted to the Michaelis-Menten equation. Mean values and standard deviations for kinetic parameters were determined from triplicate technical repeats.

#### Confocal Image Analysis

For confocal image analysis, Pearson’s R values for co-localization between EGFP-Parkin and MitoTracker were calculated from images of over 70 randomly selected EGFP-Parkin-positive cells collected from triplicated transfection experiments for each sample. The statistics and distribution of these values were analyzed using GraphPad Prism, with statistical significance determined via an unpaired Student’s T-test (*, p < 0.05; **, p < 0.005). Error bars represent mean ± SD.

#### FACS assay of the mitophagy activities

In FACS assay, over 15,000 mt-Keima positive cells were counted for each sample and the population of the cells undergoing mitophagy was gated as described(Lin, Chen et al. 2020). The statistics were calculated from the results of triplicated biological repeats. The statistics and distribution of these values were analyzed using Excel, with statistical significance determined via a paired Student’s T-test (*, p < 0.05; **, p < 0.01). Error bars represent mean ± SD.

#### Western blotting analysis

In Western blotting analysis, the band intensities were quantified with ImageJ. The statistics were determined from the results of triplicated biological repeats. The statistics and distribution of these values were analyzed using Excel, with statistical significance determined via a paired Student’s T-test (*, p < 0.05; **, p < 0.01). Error bars represent mean ± SD.

## Author Contributions

Conceptualization: LZM, XQ; Methodology: SW, YJ, XL, YZ, HX; Investigation: YJ, YZ, XL, SW, JX, YD, CC, XY, BL, HX, SC, ZL; Visualization: YZ, HX, XL, XQ, LZM; Supervision: SFS, ZL, XQ, LZM; Writing—original draft: LZM, XQ, HX, XL, YZ, ZL; Writing—review & editing: LZM, XQ, HX, XL, YZ, ZL

## Acknowledgments

The authors are grateful to Dongqing Li and Yiying Xing (Tianjin University) as well as all staff members of the Cryo-EM center (Southern University of Science and Technology) for their technical support. LZM and XQ express deep gratitude to Drs. Yufeng Yang (Fuzhou University) and Xinping Yang (Southern Medical University) for their critical reading of this manuscript. The authors are grateful to the financial support from National Natural Science Foundation of China to LZM (#31670738, #31470730), XQ (#31400645) and ZL (#82070329).

