## Supplemental Information for "DYNAMIC CONFORMATIONS IN THE ACTIVATION OF PINK1 KINASE"

#### Supplementary Information

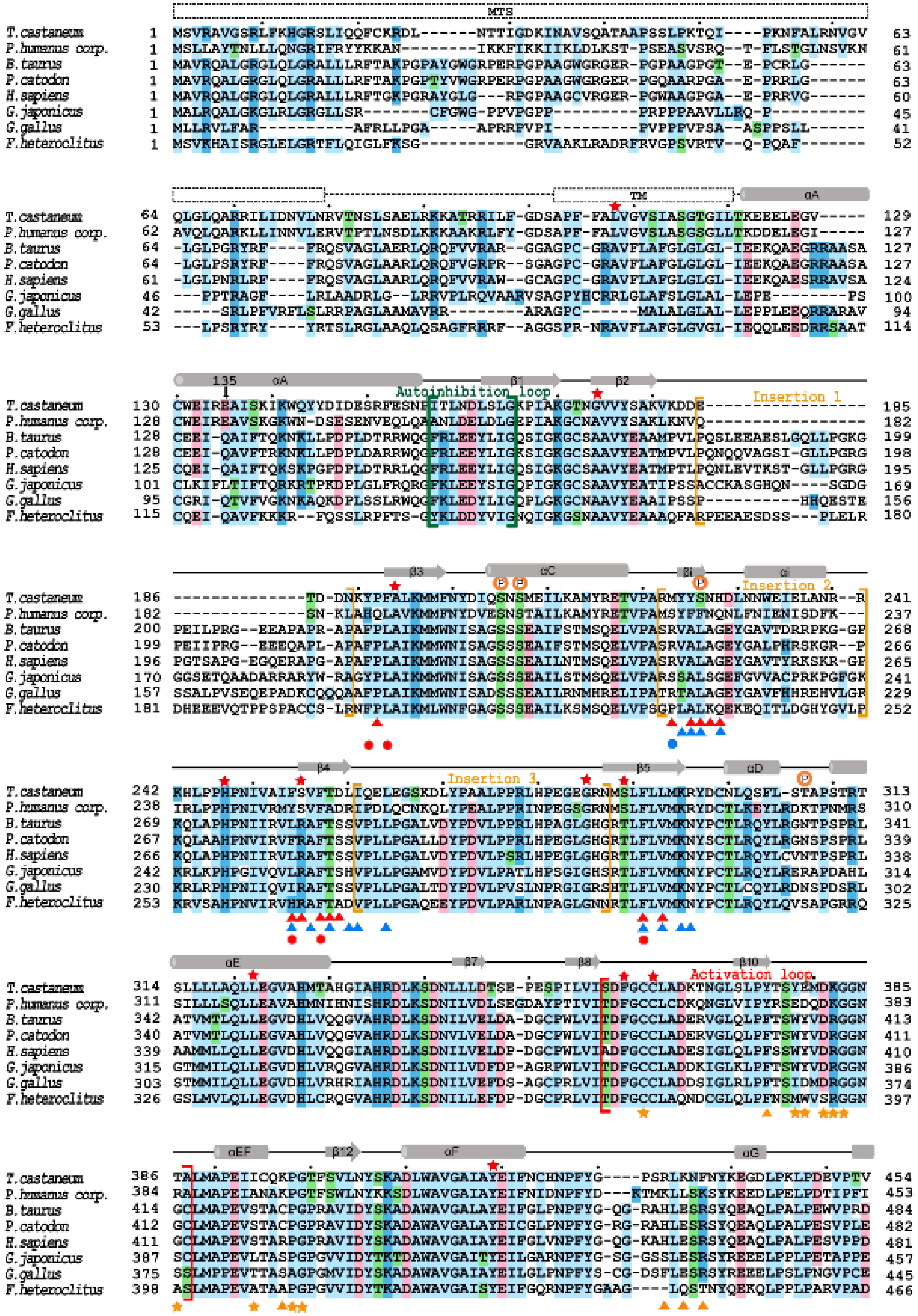

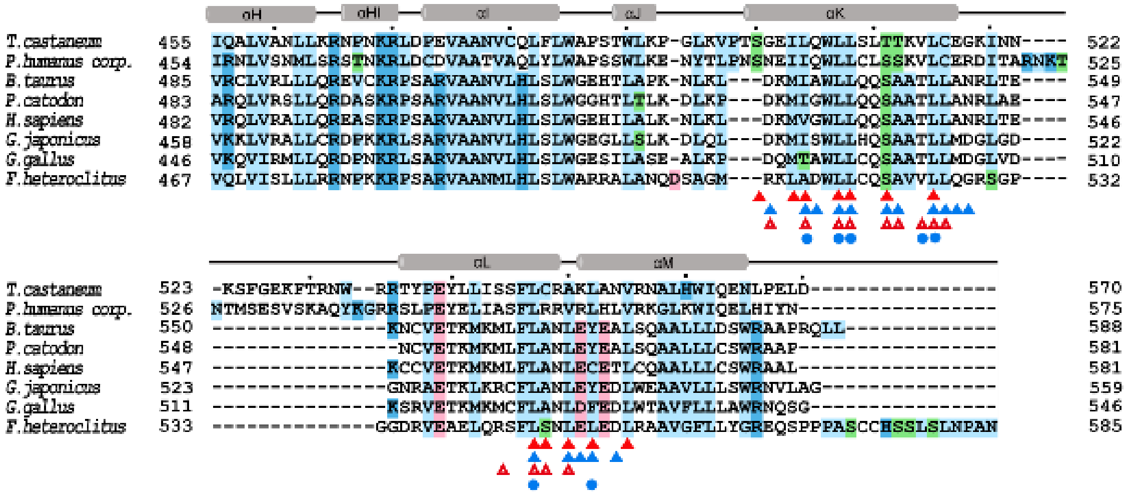

**Fig. S1. Multiple Sequence Alignment of PINK1s.** Conserved residues are highlighted with solid shades. Every 10th residue is marked with a black dot above the sequences. The α helices and β strands of PINK1 are represented by grey rods and arrows, respectively. EOPD mutations are indicated with red stars above the sequences. Known phosphorylation sites are marked with circled "P" symbols. Sequences for the three kinase insertions are enclosed in golden brackets, while the autoinhibition loop and activation loop are enclosed in green and red brackets, respectively. Residues at the dimer interface of *Tc*PINK1$^{L552R}$ are marked with solid red triangles at the bottom of the sequences, while those at the dimer interface of *Tc*PINK1$^{DDEE}$ are marked with blue triangles. Mutations that reduce Ub phosphorylation activity of *Tc*PINK1 are indicated with solid red circles, whereas mutations that enhance activity are marked with solid blue circles. Residues involved in dimeric crystallographic packing are denoted with red empty triangles. On the dimer-dimer trans-autophosphorylation interface, residues on the side of the 'kinase dimer' are marked with golden stars, while those on the side of the 'substrate dimer' are marked with golden triangles.

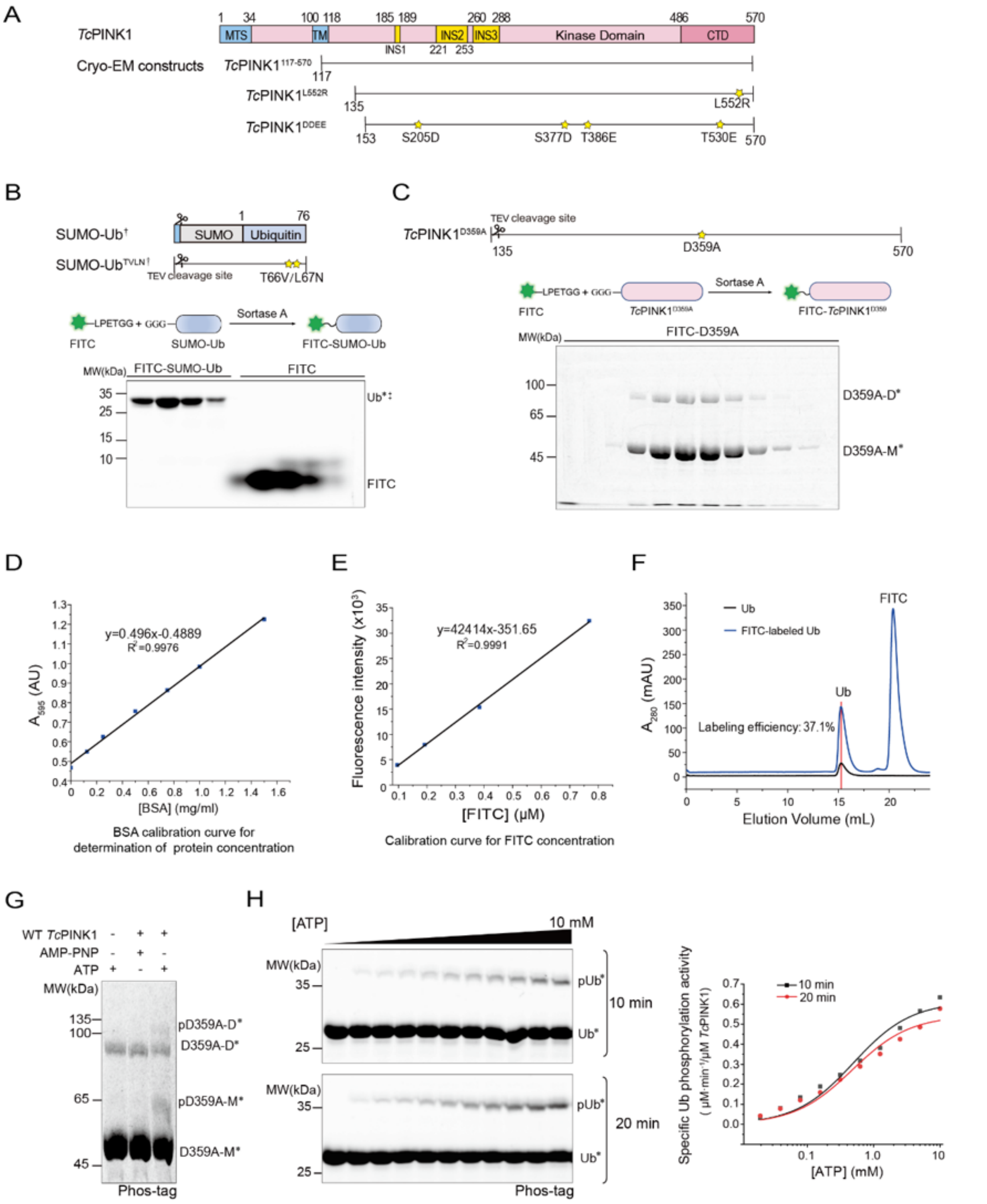


**Fig. S2. Development of a Highly Sensitive, Quantitative PINK1 Kinase Activity Assay.**
**A)** Diagram of the full-length *Tc*PINK1 sequence (top) and constructs for Cryo-EM studies (middle to bottom). Boundaries for the mitochondrion-targeting sequence (MTS), transmembrane domain (TM), insertions (INS), kinase domain, and C-terminal domain (CTD) are labeled. Mutations mimicking phosphorylated Ser/Thr residues in the *Tc*PINK1$^{DDEE}$ construct are marked with yellow stars. **B)** Fluorescent labeling of purified SUMO-Ub. Constructs for expressing and purifying SUMO-Ub and SUMO-Ub$^{TVLN}$ are shown (top). Mutated residues in SUMO-Ub$^{TVLN}$ are indicated with yellow stars. Purified SUMO-Ub with an N-terminal GGG sequence is labeled with FITC-

tagged LPETGG peptide using Sortase A (middle). The fluorescently labeled SUMO-Ub is separated from the FITC-tagged LPETGG peptide by gel-filtration chromatography (Superdex G200 column). SDS-PAGE analysis of fractionated samples reveals fluorescent bands for FITC-labeled SUMO-Ub and free FITC-peptide, visualized with a GE LAS-4000 imager (bottom). For clarity, SUMO-Ub is denoted as Ub, and SUMO-$Ub^{TVLN}$ is denoted as $Ub^{TVLN}$. FITC-labeled substrates are marked with black stars in subsequent figures. **C)** Fluorescent labeling of *Tc*$PINK1^{D359A}$ for trans-autophosphorylation assays. The construct used for expressing and purifying *Tc*$PINK1^{D359A}$ is shown (top), with the mutation marked by a yellow star. Affinity-purified *Tc*$PINK1^{D359A}$ is digested with TEV protease to expose the N-terminal Sortase recognition sequence (GGG). The digested protein is labeled with FITC-LPETGG peptide via Sortase A-mediated ligation (middle). The FITC-labeled *Tc*$PINK1^{D359A}$ is purified from the ligation mixture using a Hitrap Desalting column and analyzed by SDS-PAGE with detection by a GE LAS-4000 imager. **D)** Calibration curve for determining protein concentration. The Bradford assay, using BSA as a standard, is employed to measure protein concentrations for the calibration curve. **E)** Calibration curve for determining FITC concentration. Fluorescent intensities of FITC-LPETGG peptide are measured at 495 nm excitation and 515 nm emission, plotted against peptide concentrations. **F)** Gel-filtration chromatography profiles for the purification of FITC-labeled and non-labeled Ubs. Both labeled and non-labeled Ubs elute from the Superdex G200 column at the same position, exhibiting similar monodispersed profiles. **G)** The kinase-deficient mutant *Tc*$PINK1^{D359A}$ is specifically phosphorylated by WT *Tc*PINK1. The phosphorylation of FITC-labeled *Tc*$PINK1^{D359A}$ is compared in the absence of WT *Tc*$PINK1^{135\text{-}570}$, in the presence of WT *Tc*$PINK1^{135\text{-}570}$ with 2 μM AMP-PNP, or with 2 μM ATP. Phosphorylation is detected by fluorescent Phos-tag SDS-PAGE. **H)** Kinetics of Ub phosphorylation by *Tc*$PINK1^{135\text{-}570}$. Ub phosphorylation activity is measured as a function of ATP concentration ranging from 19.5 μM to 10 mM. After 10 or 20 minutes of reaction, phosphorylated samples are analyzed by fluorescent Phos-tag SDS-PAGE (left), and reaction kinetic parameters are determined by fitting the data to the Michaelis-Menten equation (right).

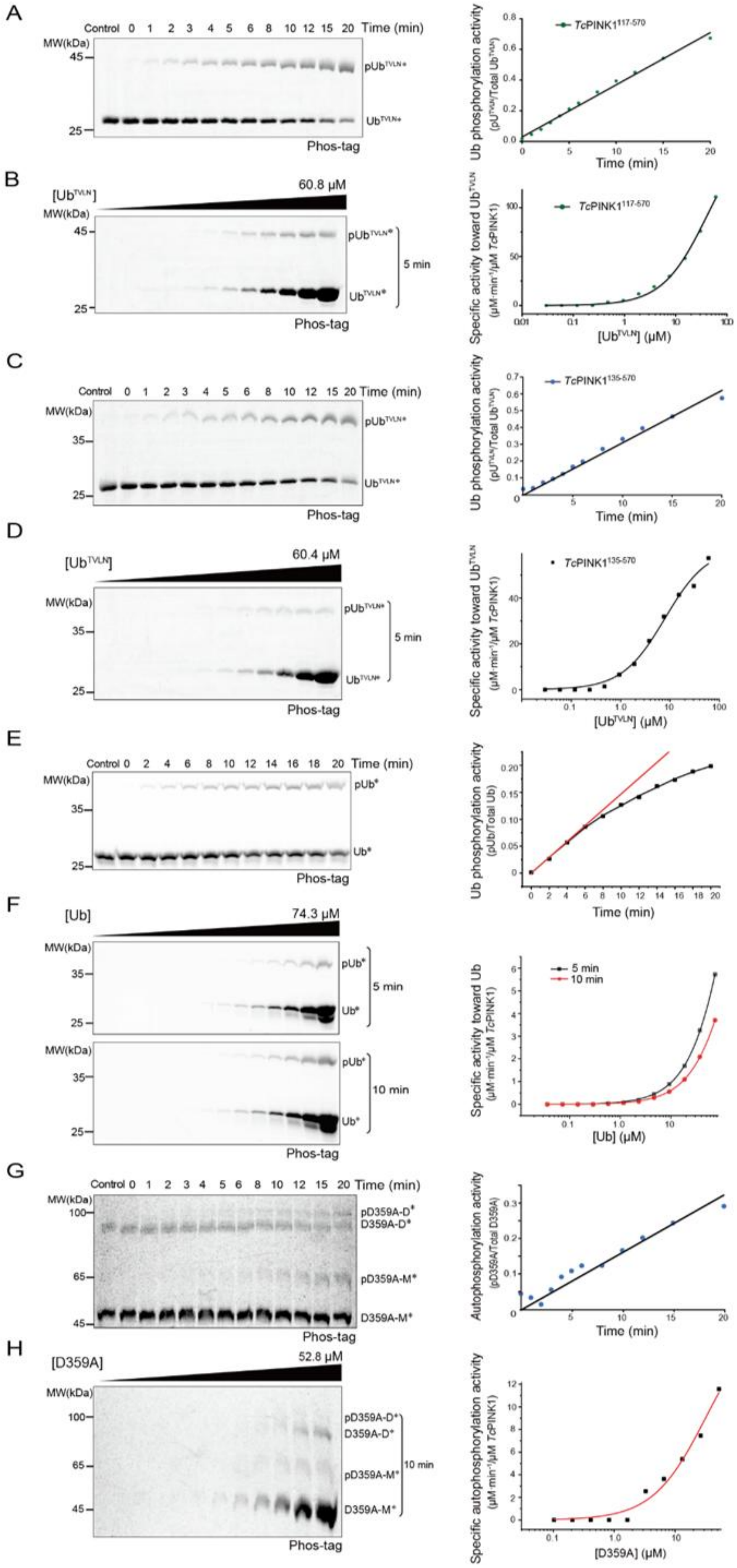
A
Control 0 1 2 3 4 5 6 8 10 12 15 20 Time (min)
MW(kDa)
45
25
pUbTVLN*
UbTVLN*
Phos-tag
TcPINK1117-570
Ub phosphorylation activity
(pUbTVLN/Total UbTVLN)
Time (min)
B
[UbTVLN]
60.8 μM
5 min
Specific activity toward UbTVLN
(μM min-1/μM TcPINK1)
[UbTVLN] (μM)
C
Control 0 1 2 3 4 5 6 8 10 12 15 20 Time (min)
MW(kDa)
35
25
TcPINK1135-570
D
[UbTVLN]
60.4 μM
E
Control 0 2 4 6 8 10 12 14 16 18 20 Time (min)
pUb*
Ub*
(pUb/Total Ub)
F
[Ub]
74.3 μM
10 min
Specific activity toward Ub
[Ub] (μM)
G
Control 0 1 2 3 4 5 6 8 10 12 15 20 Time (min)
100
65
45
pD359A-D*
D359A-D*
pD359A-M*
D359A-M*
Autophosphorylation activity
(pD359A/Total D359A)
H
[D359A]
52.8 μM
Specific autophosphorylation activity
(μM min-1/μM TcPINK1)
[D359A] (μM)

**Fig. S3. Kinetic Study of *Tc*PINK1 Kinase Activities. A)** Reaction rate of the phosphorylation of $Ub^{TVLN}$ mutant by *Tc*$PINK1^{117\text{-}570}$. 10.0 μM FITC-labeled $Ub^{TVLN}$ was phosphorylated by 0.01 μM *Tc*$PINK1^{117\text{-}570}$. Reactions were stopped at indicated time points, and phosphorylated samples were analyzed. The reaction rate remains linear for up to 20 minutes. **B)** Kinetics in the phosphorylation of $Ub^{TVLN}$ by *Tc*$PINK1^{117\text{-}570}$. FITC-labeled $Ub^{TVLN}$ at concentrations ranging from 0.03 μM to 60.8 μM was phosphorylated by 0.01 μM WT *Tc*$PINK1^{117\text{-}570}$. After 5 minutes, samples were separated by Phos-tag SDS-PAGE for analysis. **C)** Reaction rate of the phosphorylation of $Ub^{TVLN}$ mutant by *Tc*$PINK1^{135\text{-}570}$. 8.0 μM FITC-labeled $Ub^{TVLN}$ was phosphorylated by 0.01 μM *Tc*$PINK1^{135\text{-}570}$. Reactions were stopped at indicated time points for analysis. The reaction rate remains linear for up to 20 minutes. **D)** Kinetics in phosphorylation activities of $Ub^{TVLN}$ by *Tc*$PINK1^{135\text{-}570}$. FITC-labeled $Ub^{TVLN}$ at concentrations ranging from 0.029 μM to 60.4 μM was phosphorylated by 0.01 μM WT *Tc*$PINK1^{135\text{-}570}$. After 5 minutes, samples were analyzed by Phos-tag SDS-PAGE. **E)** Reaction rate of the phosphorylation of Ub by *Tc*$PINK1^{135\text{-}570}$. 7.46 μM FITC-labeled Ub was phosphorylated by 0.3 μM $TcPINK1^{135\text{-}570}$. Reactions were stopped at indicated time points, and samples were analyzed by fluorescent Phos-tag SDS-PAGE (left). The initial reaction rate was determined from the tangent line (red) of the reaction curve (black) at 0 minutes; the curve deviated from the tangent line after 10 minutes. **F)** Kinetics in phosphorylation of Ub by *Tc*$PINK1^{135\text{-}570}$. Purified FITC-labeled Ub at concentrations ranging from 0.036 μM to 74.3 μM was phosphorylated by 0.3 μM WT *Tc*$PINK1^{135\text{-}570}$. Samples were analyzed by fluorescent Phos-tag SDS-PAGE after 5 or 10 minutes (left), and kinetic data were fitted to the Michaelis-Menten equation (right). **G)** Reaction rate of the phosphorylation of *Tc*$PINK1^{D359A}$ mutant by WT *Tc*$PINK1^{135\text{-}570}$. 13.9 μM FITC-labeled *Tc*$PINK1^{D359A}$ was phosphorylated by 0.03 μM WT *Tc*$PINK1^{135\text{-}570}$. Reactions were stopped at indicated time points for analysis, with constant reaction rates observed over 20 minutes. **H)** Kinetics in trans-autophosphorylation of *Tc*$PINK1^{D359A}$ by *Tc*$PINK1^{135\text{-}570}$. FITC-labeled *Tc*$PINK1^{D359A}$ at concentrations ranging from 0.10 μM to 52.8 μM was phosphorylated by 0.03 μM WT *Tc*$PINK1^{135\text{-}570}$. Reactions were stopped after 10 minutes, and samples were analyzed as described.

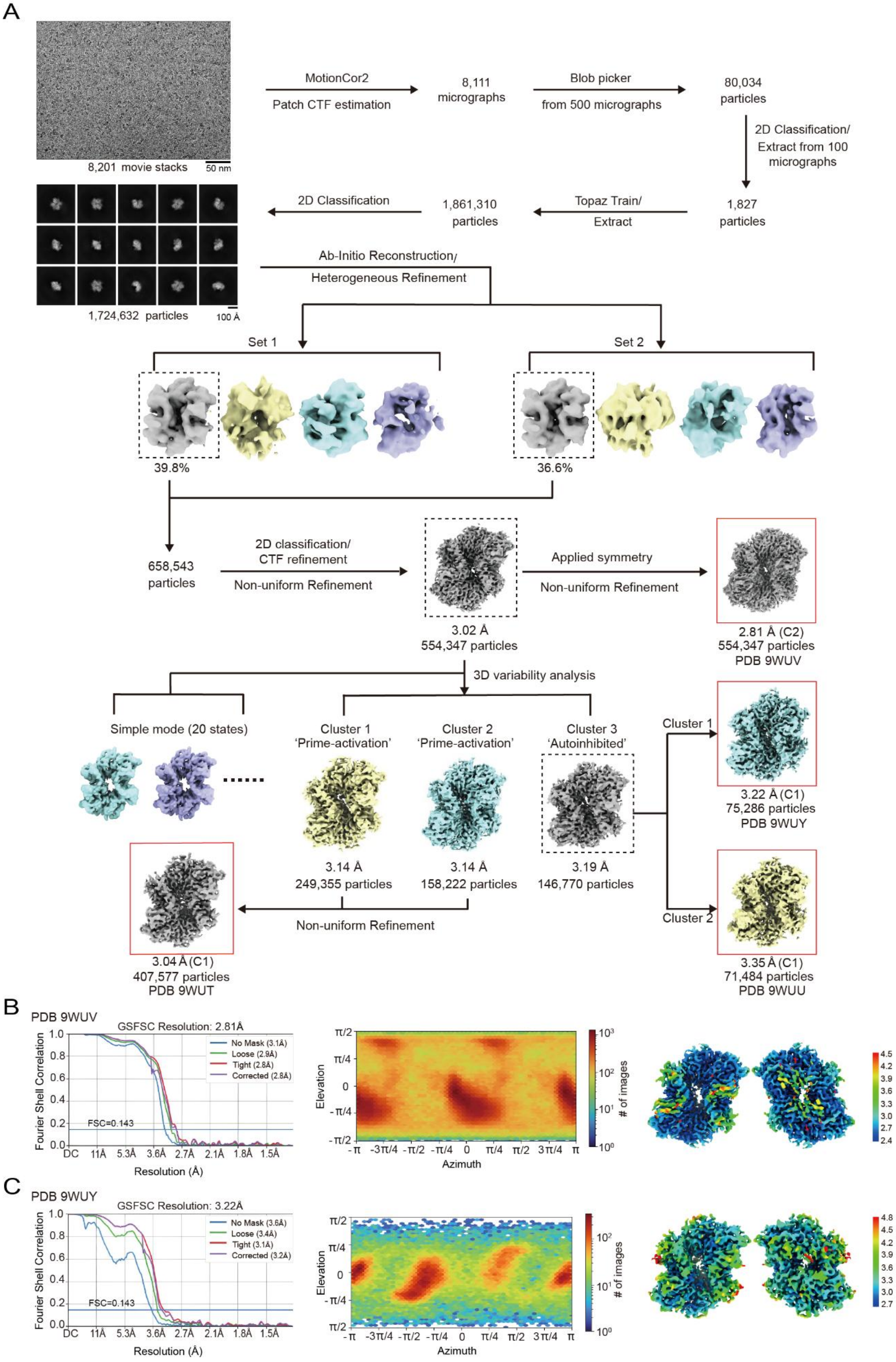

A
8,201 movie stacks
50 nm
MotionCor2
Patch CTF estimation
8,111 micrographs
Blob picker
from 500 micrographs
80,034 particles
2D Classification/ Extract from 100 micrographs
1,827 particles
Topaz Train/ Extract
1,861,310 particles
2D Classification
1,724,632 particles
100 Å
Ab-Initio Reconstruction/
Heterogeneous Refinement
Set 1
Set 2
39.8%
36.6%
658,543 particles
2D classification/ CTF refinement
Non-uniform Refinement
3.02 Å
554,347 particles
Applied symmetry
Non-uniform Refinement
2.81 Å (C2)
554,347 particles
PDB 9WUV
3D variability analysis
Simple mode (20 states)
Cluster 1 'Prime-activation'
Cluster 2 'Prime-activation'
Cluster 3 'Autoinhibited'
3.14 Å
249,355 particles
3.14 Å
158,222 particles
3.19 Å
146,770 particles
Non-uniform Refinement
3.04 Å (C1)
407,577 particles
PDB 9WUT
Cluster 1
3.22 Å (C1)
75,286 particles
PDB 9WUY
Cluster 2
3.35 Å (C1)
71,484 particles
PDB 9WUU
B
PDB 9WUV
GSFSC Resolution: 2.81Å
Fourier Shell Correlation
No Mask (3.1Å)
Loose (2.9Å)
Tight (2.8Å)
Corrected (2.8Å)
FSC=0.143
DC 11Å 5.3Å 3.6Å 2.7Å 2.1Å 1.8Å 1.5Å
Resolution (Å)
Elevation
Azimuth
### of images
C
PDB 9WUY
GSFSC Resolution: 3.22Å
Fourier Shell Correlation
No Mask (3.6Å)
Loose (3.4Å)
Tight (3.1Å)
Corrected (3.2Å)
FSC=0.143
DC 11Å 5.3Å 3.6Å 2.7Å 2.1Å 1.8Å 1.5Å
Resolution (Å)
Elevation
Azimuth
### of images

**Fig. S4. Overall Strategies and Statistics in Processing Cryo-EM Data of *Tc*PINK1$^{L552R}$.**
**A)** Overview of the strategies used in Cryo-EM data processing. Micrographs of the *Tc*PINK1$^{L552R}$ sample were captured at a magnification of 130,000 using an FEI Titan Krios, with a pixel size of 0.668 Å/pixel and a defocus range of -1.2 μm to -1.8 μm. Particles were extracted using Topaz Train and subjected to 2D classification and 3D reconstruction with CryoSPARC. Density maps used for model building and structural analysis are highlighted with red boxes.
**B)** Statistics for the density maps corresponding to PDB 8GNC. The Fourier Shell Correlation (FSC) between the two half-density maps is shown, calculated with and without masks (left). Resolutions were determined using the gold-standard FSC 0.143 criterion. The angular distribution of particle images used to construct the density map is presented (middle). Local resolutions of the Cryo-EM density map, determined using ResMap, are shown with pseudo-colors (right). **C)** Statistics for the density maps corresponding to PDB 8ATQ. The Fourier shell correlation between the two half-density maps is presented (left), along with resolutions determined using the gold-standard FSC 0.143 criterion. The angular distribution of particle images used for the density map is shown (middle), and local resolutions for the Cryo-EM density map are displayed with pseudo-colors (right).

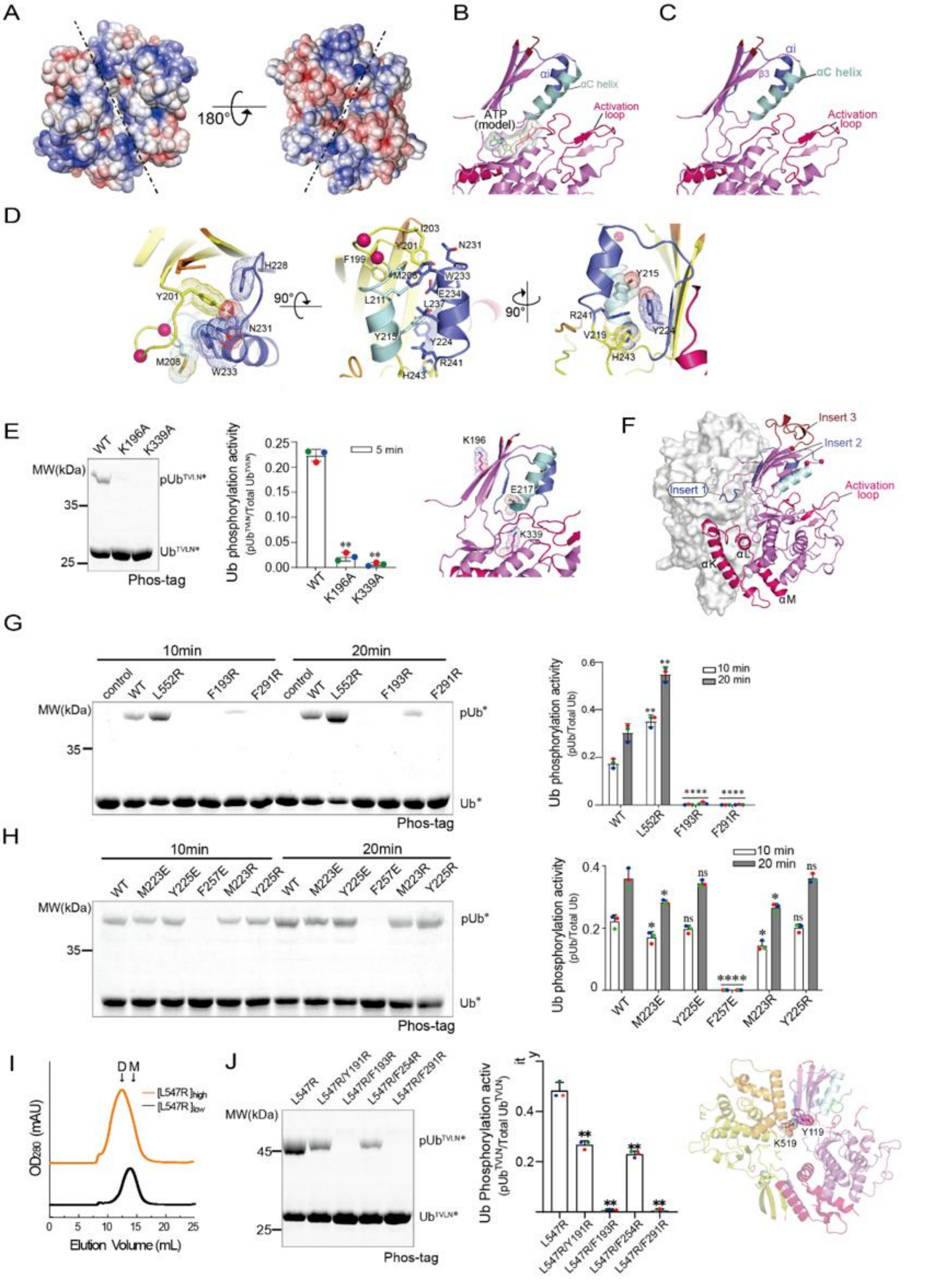


**Fig. S5. Cryo-EM Structure of Dimeric *Tc*PINK1$^{L552R}$ in the Conformation Priming for Activation. A)** Electrostatic potential on the surface of *Tc*PINK1$^{L552R}$ priming for activation. The surface of the dimer exhibits bipolar charge distribution. **B, C)** Model of ATP binding to the active site of *Tc*PINK1$^{L552R}$. Bound ATP is represented as green sticks and masked with surface representation. The model was generated by superimposing the ANP-bound *Tc*PINK1 structure (PDB ID 5YJ9) onto one subunit. **D)** Extensive interactions between the αC and αi helices in the *Tc*PINK1$^{L552R}$ structure. The αC and αi helices interact extensively along their lengths (middle). At the N-termini, residues H228, Y201, W233, N231, and M208 form a hydrophobic sandwich (left). In the middle, residues Y215, Y224, and L237 contribute hydrophobic interactions (middle). At the C-termini, R241 fits into an aromatic cage formed by Y215, Y224, and H243, creating π-cation interactions (right). Key residues are shown in stick representation. **E)** Comparison of K339A and K196A mutations on the phosphorylation of Ub$^{TVLN}$. The K196A mutation does not completely abolish the kinase activity of *Tc*PINK1, with experiments performed as described in panel D. These two residues are highlighted in the structure with sticks. **F)** Insertion 1 of the kinase is identified at the dimer interface in the low-resolution structure of *Tc*PINK1$^{L552R}$. The low-resolution density map was utilized for the refinement of the *Tc*PINK1$^{L552R}$ structure using Rosetta. One kinase subunit is shown in surface representation, while the other is depicted in cartoon. The three kinase insertions, activation loop, and α helices of the CTD are labeled. **G, H)** Effect of *Tc*PINK1$^{L552R}$ dimerization on Ub phosphorylation activity. Several hydrophobic residues at the dimer interface were mutated to charged residues to disrupt dimerization. The Ub phosphorylation activities of these mutants were compared to the WT kinase using the fluorescent Phos-tag SDS-PAGE assay (left), with statistical analysis based on triplicate technical repeats (right). **I)** Affinity-purified *Tc*PINK1$^{L547R}$ at high (~18 mg/mL) and low concentrations (~6 mg/mL) were fractionated using a Superdex G200 column. D, dimers; M, monomers. **J)** Mutations at the dimer interface of *Tc*PINK1$^{L552R}$ were introduced in *Tc*PINK1$^{L547R}$. The activities of these double mutants on phosphorylation of Ub$^{TVLN}$ were compared to *Tc*PINK1$^{L547R}$ using the fluorescent Phos-tag SDS-PAGE assay (left), with statistical analysis based on triplicate technical repeats (middle). These mutated residues are highlighted in the structure with sticks (right).

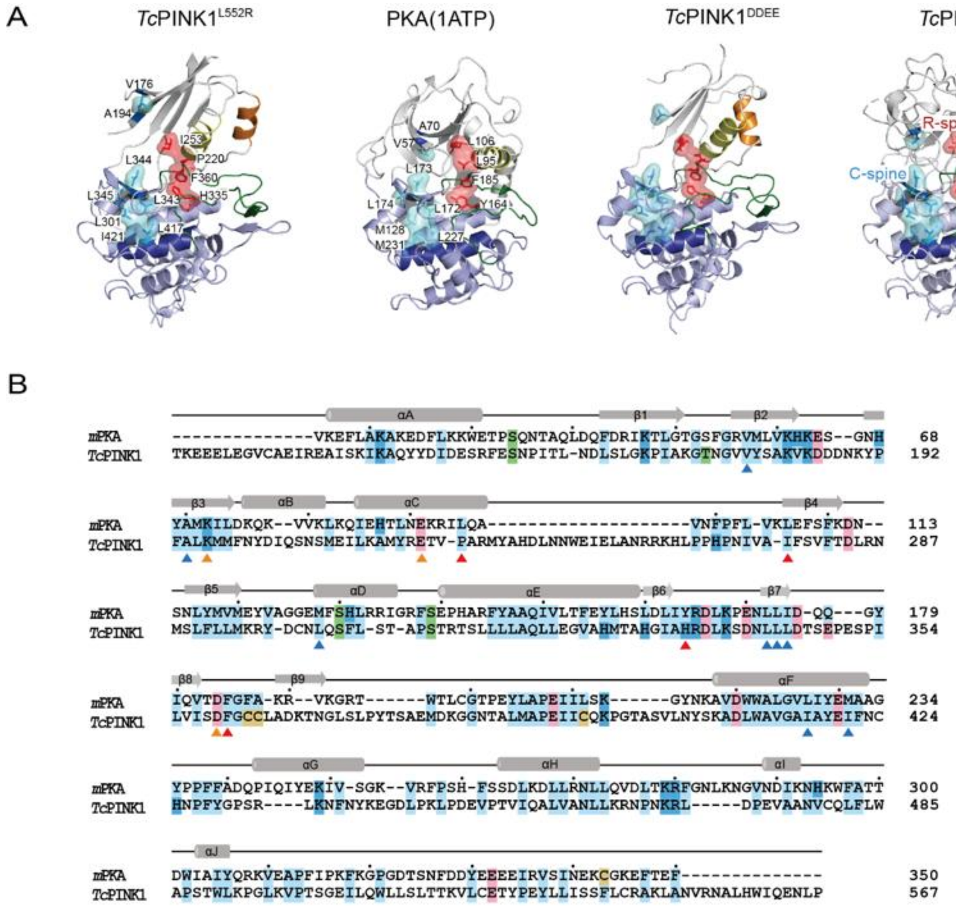


**Fig. S6. Structural Comparison of Different *Tc*PINK1 Conformations with the Active PKA Structure. A)** Comparison of various *Tc*PINK1 structures with the active PKA structure. *Tc*PINK1 and PKA structures are represented as cartoons. R-spine residues are shown as red sticks with a transparent surface mask, while C-spine residues are depicted as cyan sticks under transparent masks. **B)** Sequence alignment of *Tc*PINK1 (PDB 7MP8) and PKA (1ATP) using Secondary Structure Matching. Conserved residues are highlighted with solid shades. Every 10th residue in the PKA sequence is marked with a black dot. α helices and β strands are represented with grey rods and arrows, respectively. R-spine residues are indicated with red triangles, C-spine residues with blue triangles, and the essential catalytic triad with yellow triangles.

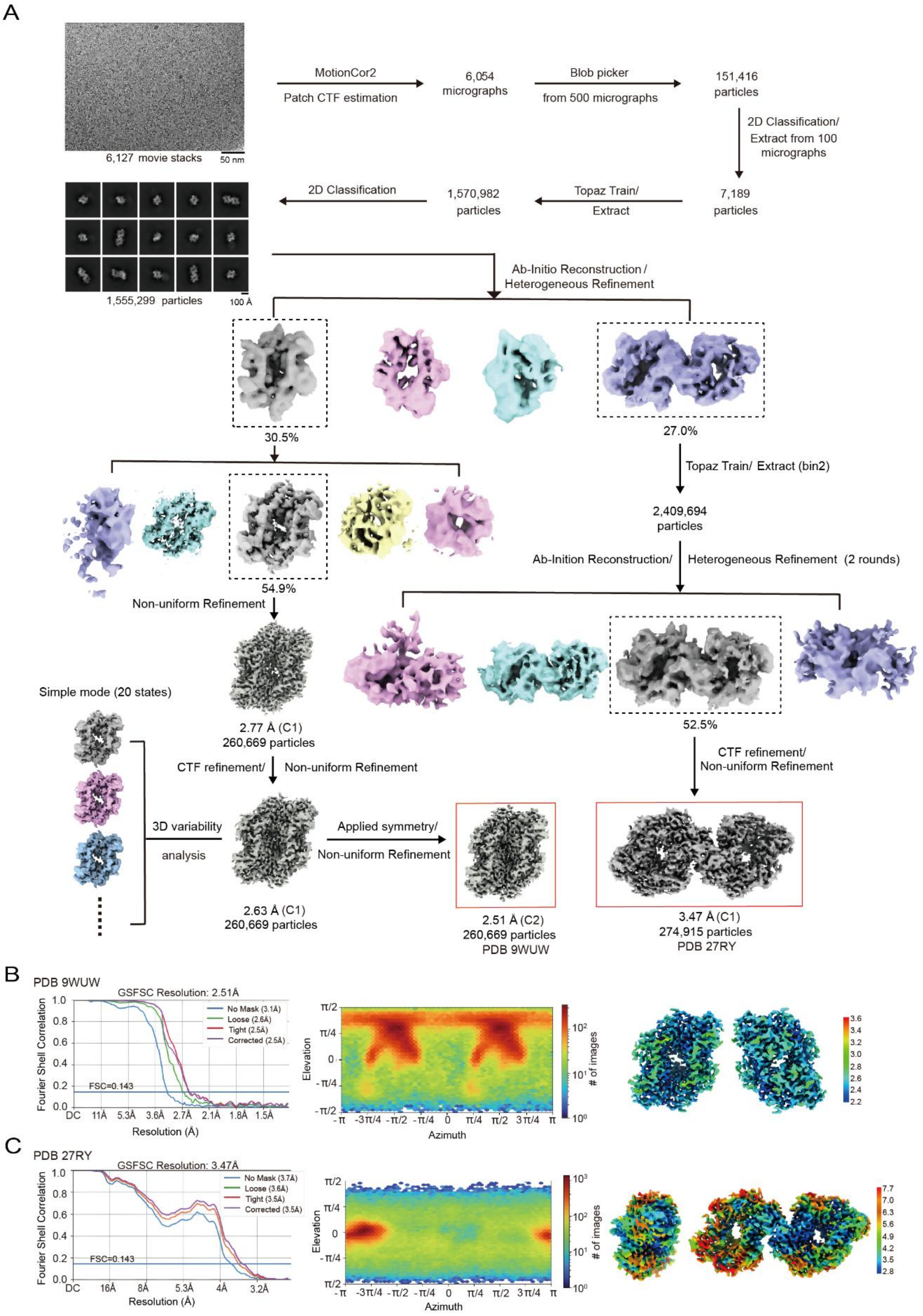

A
6,127 movie stacks
50 nm
MotionCor2
Patch CTF estimation
6,054 micrographs
Blob picker
from 500 micrographs
151,416 particles
2D Classification/ Extract from 100 micrographs
7,189 particles
Topaz Train/ Extract
1,570,982 particles
2D Classification
1,555,299 particles
100 Å
Ab-Initio Reconstruction / Heterogeneous Refinement
30.5%
27.0%
Topaz Train/ Extract (bin2)
2,409,694 particles
Ab-Inition Reconstruction/ Heterogeneous Refinement (2 rounds)
54.9%
Non-uniform Refinement
52.5%
Simple mode (20 states)
2.77 Å (C1)
260,669 particles
CTF refinement/ Non-uniform Refinement
CTF refinement/ Non-uniform Refinement
3D variability analysis
Applied symmetry/ Non-uniform Refinement
2.63 Å (C1)
260,669 particles
2.51 Å (C2)
260,669 particles
PDB 9WUW
3.47 Å (C1)
274,915 particles
PDB 27RY
B
PDB 9WUW
GSFSC Resolution: 2.51Å
No Mask (3.1Å)
Loose (2.6Å)
Tight (2.5Å)
Corrected (2.5Å)
FSC=0.143
Fourier Shell Correlation
Resolution (Å)
Elevation
Azimuth
### of images
C
PDB 27RY
GSFSC Resolution: 3.47Å
No Mask (3.7Å)
Loose (3.6Å)
Tight (3.5Å)
Corrected (3.5Å)
FSC=0.143
Fourier Shell Correlation
Resolution (Å)
Elevation
Azimuth
### of images

**Fig. S7. Overall Strategies and Statistics in Processing Cryo-EM Data of *Tc*PINK1$^{DDEE}$.**
**A)** Overview of strategies for processing Cryo-EM data to determine the dimeric and tetrameric structures of *Tc*PINK1$^{DDEE}$. Micrographs of the *Tc*PINK1$^{DDEE}$ sample were captured at a magnification of 130,000 using an FEI Titan Krios, with a pixel size of 0.668 Å/pixel and a defocus range of -1.2 μm to -1.8 μm. Particles were extracted using Topaz Train and subjected to 2D classification and 3D reconstruction with CryoSPARC. Density maps used for model building and structural analysis are highlighted with red boxes. **B)** Statistics for the density maps of dimeric *Tc*PINK1$^{DDEE}$ (PDB 8GNM). The FSC between the two half-density maps is shown (left), calculated with and without masks. Resolutions were determined using the gold-standard FSC criterion. The angular distribution of particle images used for constructing the density map is presented (middle). Local resolutions of the Cryo-EM density map for dimeric *Tc*PINK1$^{DDEE}$ are displayed with pseudo-colors (right). **C)** Statistics for the density maps of tetrameric *Tc*PINK1$^{DDEE}$ (PDB 8GO4). The FSC between the two half-density maps is shown (left), with FSC curves calculated with and without masks. Resolutions were determined using the gold-standard criterion. The angular distribution of particle images used for constructing the density map of tetrameric *Tc*PINK1$^{DDEE}$ is shown (middle), noting that preferred orientation led to anisotropic resolution. Local resolutions for the Cryo-EM density map of tetrameric *Tc*PINK1$^{DDEE}$ are represented with pseudo-colors (right).

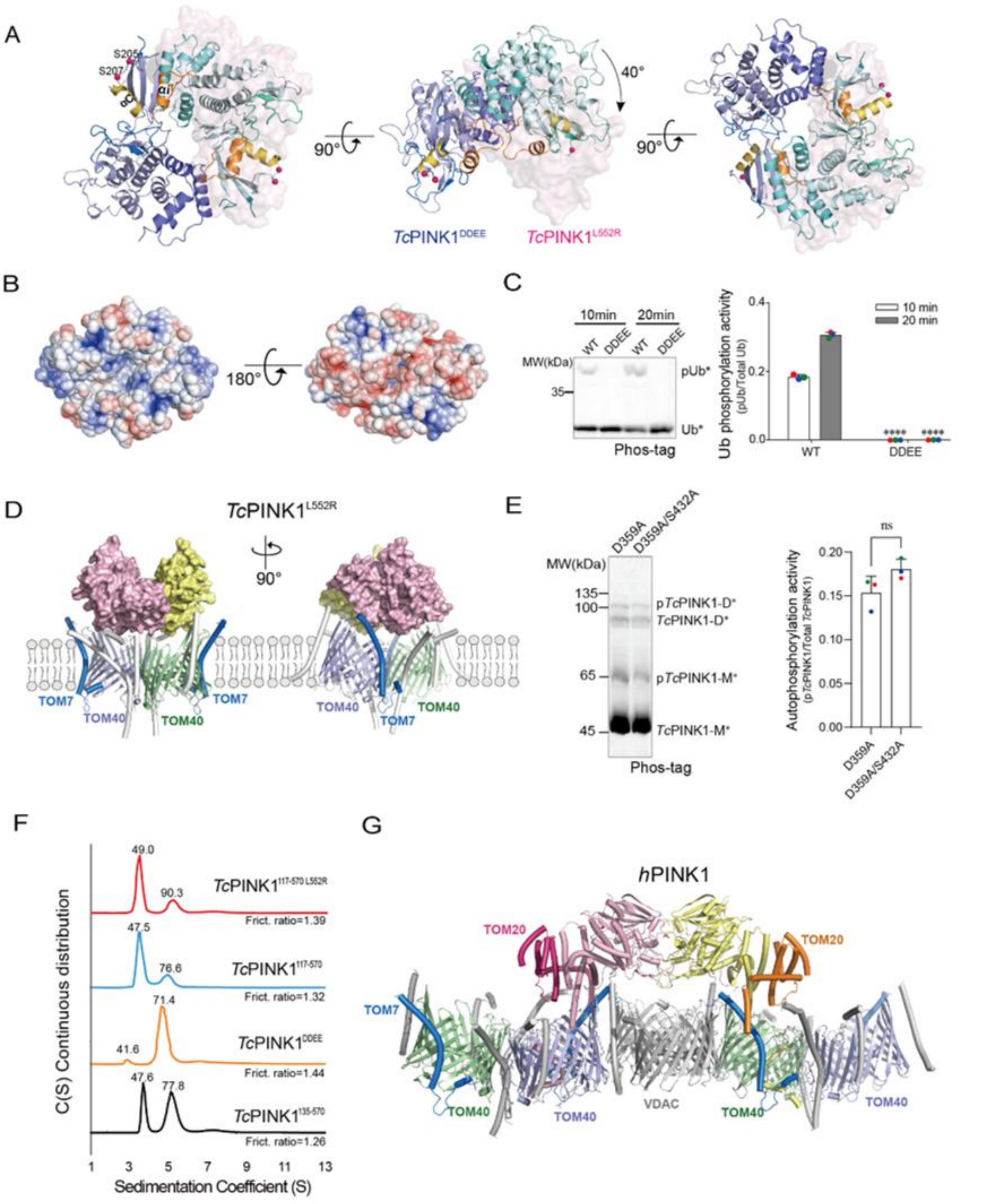


**Fig. S8. Cryo-EM Structure of the Inactive, Phosphomimetic Mutant *Tc*PINK1$^{DDEE}$.**
**A)** Comparison of dimeric structures of *Tc*PINK1$^{DDEE}$ and *Tc*PINK1$^{L552R}$. The structures were superimposed on one subunit of the dimer. The non-superimposed subunit of *Tc*PINK1$^{DDEE}$ is shown in cyan cartoon, while its counterpart in *Tc*PINK1$^{L552R}$ is depicted in pink surface. The superimposed subunit of *Tc*PINK1$^{L552R}$ is omitted for clarity. The αC is colored yellow, and insertion 2 is shown in gold. **B)** Bipolar electrostatic potential on the surface of the *Tc*PINK1$^{DDEE}$ dimer. **C)** The phosphomimetic mutant *Tc*PINK1$^{DDEE}$ exhibits reduced Ub phosphorylation activity. The kinase activities of WT *Tc*PINK1 and the *Tc*PINK1$^{DDEE}$ mutant were

assessed using the fluorescent Phos-tag SDS-PAGE assay (left), with statistical analysis of triplicate technical repeats (right). **D)** Docking studies of *Tc*PINK1 dimers onto the dimeric TOM complex were conducted, focusing on their interactions with TOM7 and shape complementarity. The structure of the *Tc*PINK1$^{L552R}$ dimer demonstrates compatibility with the dimeric TOM complex, exhibiting complementary shape and electrostatic potential. The *Tc*PINK1 dimer is represented in a cartoon under surface, while the TOM complex is depicted in a cartoon. **E)** S432 is not a major phosphorylation site for *Tc*PINK1. The mutants *Tc*PINK1$^{D359A}$ and *Tc*PINK1$^{D359A/S432A}$ were labeled with FITC and subjected to phosphorylation by *Tc*PINK1$^{135\text{-}570}$. The phosphorylation levels of *Tc*PINK1$^{D359A}$ and *Tc*PINK1$^{D359A/S432A}$ were detected by fluorescent Phos-tag SDS-PAGE, with results shown on the left and statistical analysis of triplicate technical repeats on the right. **F)** AUC analysis of purified WT *Tc*PINK1$^{117\text{-}570\ L552R}$, *Tc*PINK1$^{117\text{-}570}$, *Tc*PINK1$^{DDEE}$, and *Tc*PINK1$^{135\text{-}570}$. **G)** The structure of *h*PINK1 priming dimer associated with TOM and VDAC2 complexes. *h*PINK1s, pink and yellow rods; TOM7s, teal rods; TOM40s, green and slate cartoons; VDAC2, grey cartoons.

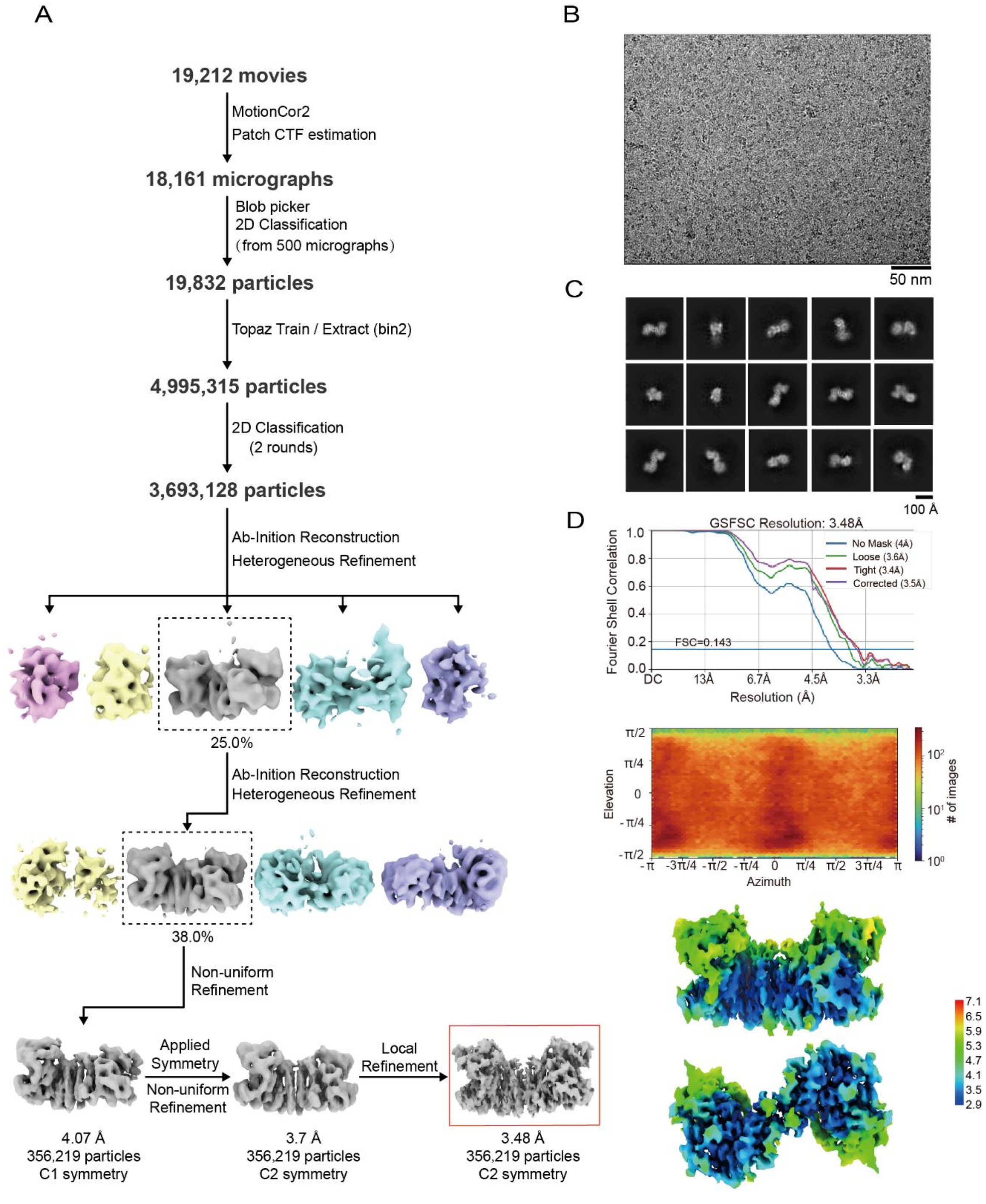


**Fig. S9. Strategies and Statistics for Processing Cryo-EM Data of *Tc*PINK1$^{117\text{-}570}$.**
**A)** Overview of the strategies used in Cryo-EM data processing. Micrographs of the *Tc*PINK1$^{117\text{-}570}$ sample were captured at a magnification of 130,000 using an FEI Titan Krios. The pixel size was 0.668 Å/pixel, and the defocus values ranged from -1.2 μm to -1.8 μm. *Tc*PINK1$^{117\text{-}570}$ particles were extracted using Topaz Train and processed through 2D classification and 3D reconstruction with CryoSPARC. The density maps used for model building and structural analysis are highlighted with a red box. **B)** Representative raw image of *Tc*PINK1$^{117\text{-}570}$. **C)** 2D

class averages of *Tc*PINK1$^{117\text{-}570}$ particles. **D)** Statistics for the model density maps. The FSC between the two half-density maps is shown (top), calculated with and without masks. Resolutions were determined using the gold-standard FSC 0.143 criterion. The angular distribution of particle images used for constructing the density map is shown (middle). Local resolutions of the Cryo-EM density map were assessed with ResMap and represented on the map using pseudo-colors (bottom).

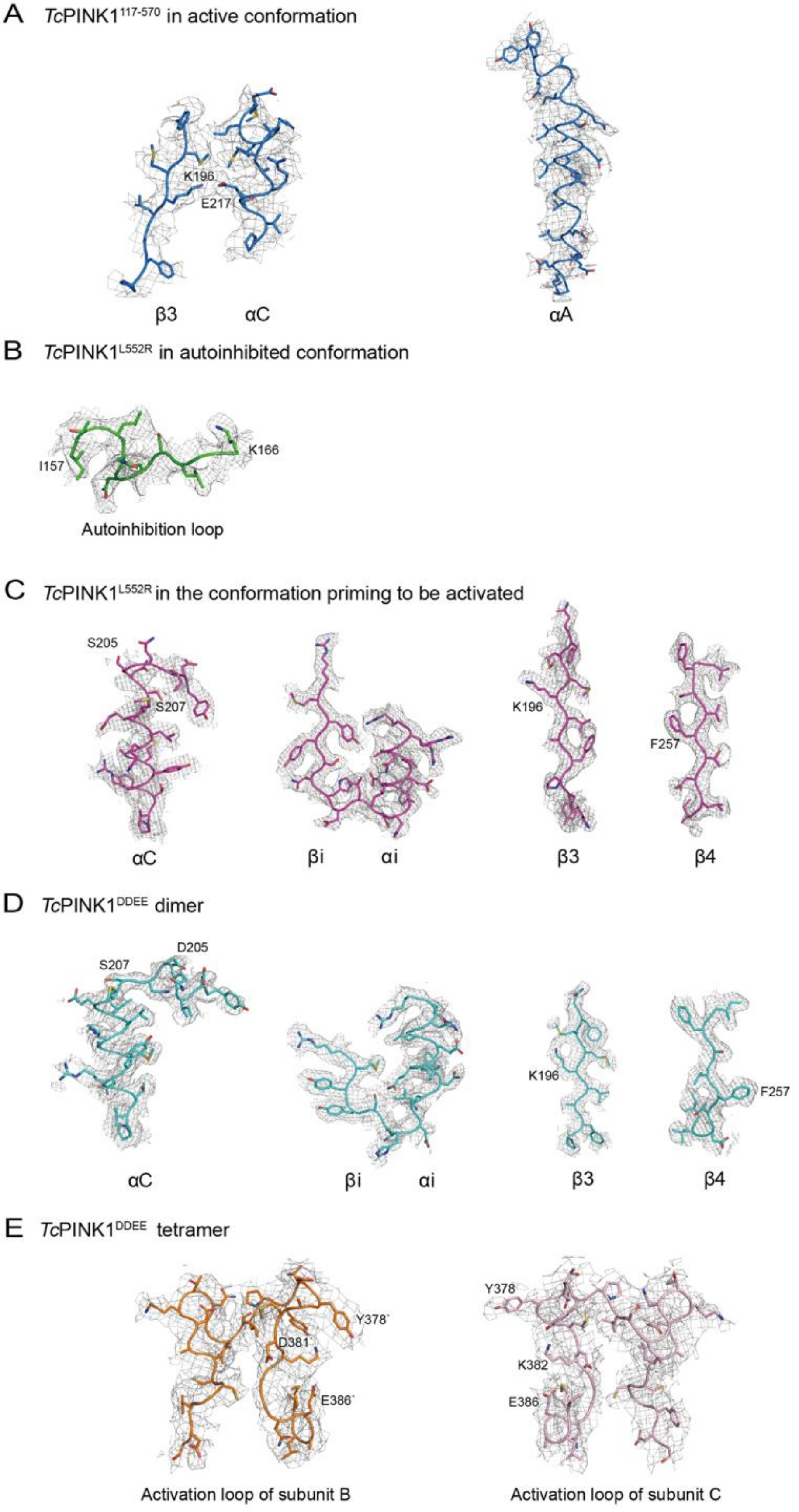
A TcPINK1117-570 in active conformation
K196
E217
β3
αC
αA
B TcPINK1L552R in autoinhibited conformation
I157
K166
Autoinhibition loop
C TcPINK1L552R in the conformation priming to be activated
S205
S207
K196
F257
αC
βi
αi
β3
β4
D TcPINK1DDEE dimer
S207
D205
K196
F257
αC
βi
αi
β3
β4
E TcPINK1DDEE tetramer
Y378`
D381`
E386`
Y378
K382
E386
Activation loop of subunit B
Activation loop of subunit C

**Fig. S10. Representative Cryo-EM Density Maps for Key Structural Elements of *Tc*PINK1.**
**A)** Cryo-EM density maps for the β3 strand, αC helix (left), and the αA helix (right) in the Cryo-EM structure of activated TcPINK1$^{117\text{-}570}$. Density maps are displayed as isomeshes contoured at the 4σ level. Main chains are shown as cartoon loops, with side chains represented as sticks.
**B)** Cryo-EM density maps for the autoinhibition loop in the Cryo-EM structure of autoinhibited *Tc*PINK1$^{L552R}$. Density maps are contoured at the 4σ level, with main chains as cartoon loops and side chains as sticks. **C)** Cryo-EM density maps for the αC helix, insertion 2 (including the αi helix and βi strand), β3, and β4 strands of *Tc*PINK1$^{L552R}$ in a primed activation structure, contoured at the 4σ level. **D)** Cryo-EM density maps for the αC helix, insertion 2, β3, and β4 strands of dimeric *Tc*PINK1$^{DDEE}$, contoured at the 4σ level. **E)** Cryo-EM density maps for the activation loops of subunit B and subunit C in the tetrameric structure of *Tc*PINK1$^{DDEE}$, contoured at the 4σ level.

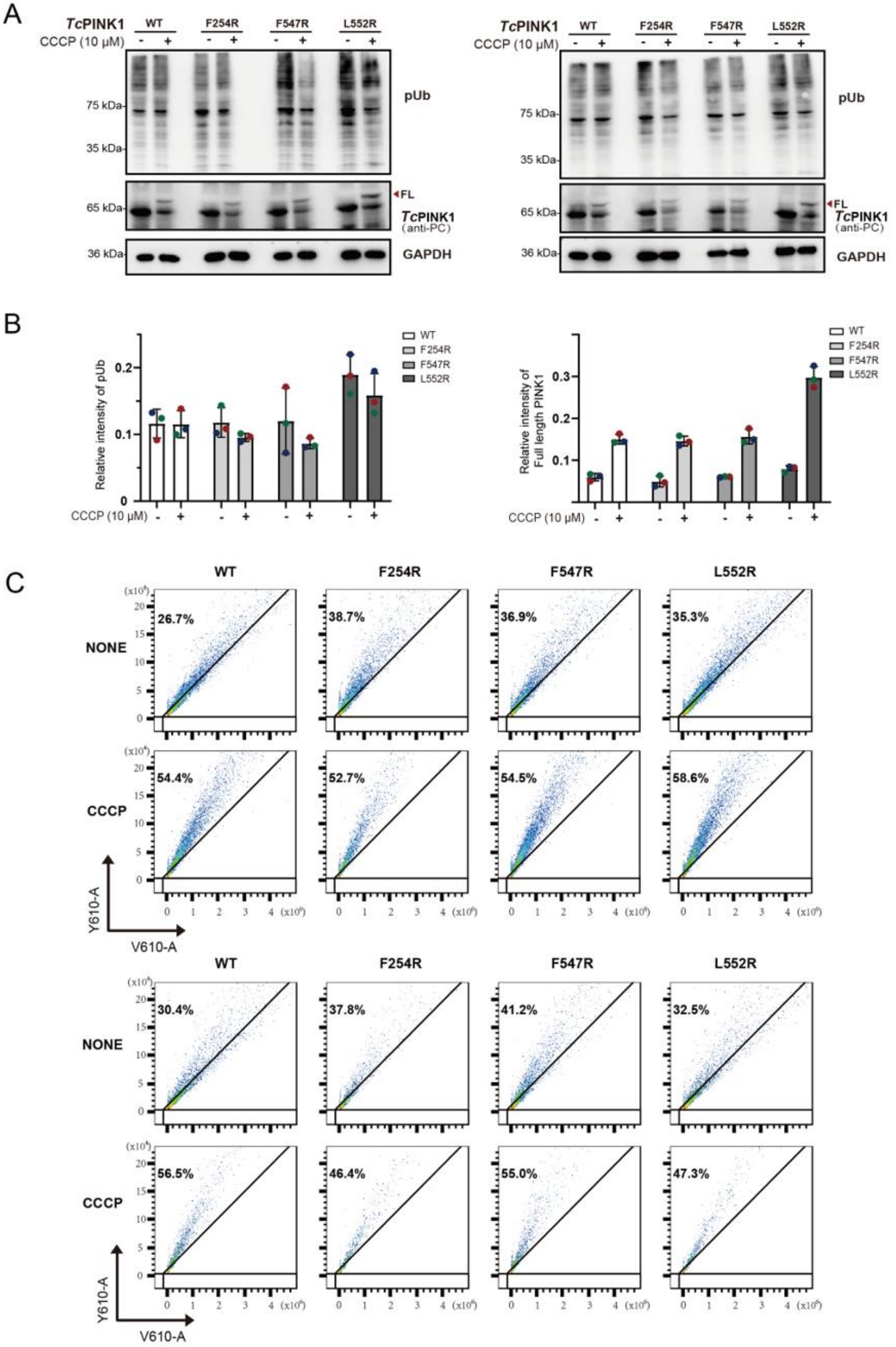
A
TcPINK1
CCCP (10 μM)
WT
F254R
F547R
L552R
pUb
75 kDa
35 kDa
65 kDa
FL
TcPINK1 (anti-PC)
36 kDa
GAPDH
B
Relative intensity of pUb
Relative intensity of Full length PINK1
C
NONE
CCCP
26.7%
38.7%
36.9%
35.3%
54.4%
52.7%
54.5%
58.6%
30.4%
37.8%
41.2%
32.5%
56.5%
46.4%
55.0%
47.3%
Y610-A
V610-A

**Fig. S11. The Expression, Phosphorylation, and Mitophagy Activities of *Tc*PINK1 WT and Mutants. A,B)** Results from two replicated biological experiments in analyzing the expression, processing, and Ub phosphorylation activities of *Tc*PINK1s transfected into *hPINK1*-KO COS7 cells. The transfected cells were treated with or without 10 μM CCCP for 8 hours before subjecting to the Western blotting analysis with antibodies against pUb, protein C, and GAPDH, respectively. The full-length *Tc*PINK1s are marked with a black arrow. **C,D)** The statistical analysis of the levels of Ub phosphorylation by *Tc*PINK1s and the accumulation of the full-length *Tc*PINK1s (D) with or without CCCP treatment. The statistics were determined from triplicate biological repeats, with error bars representing the mean ± SD. **E)** Results from two replicated biological experiments in flowcytometry analysis of the mitophagy activities of *Tc*PINK1 WT and mutants transfected in *hPINK1*-KO HeLa cells. The transfected cells were treated with or without 10 μM CCCP for 10 hours before the flowcytometry assay.

F1-D

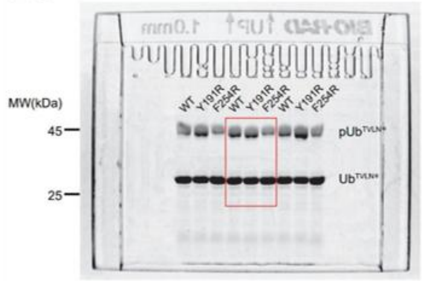


F2-D

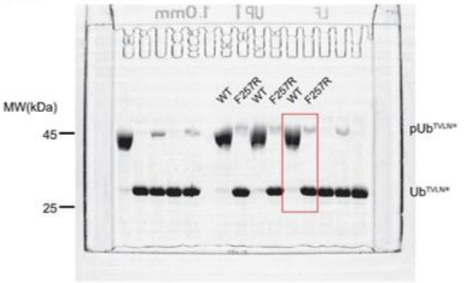


F3-F

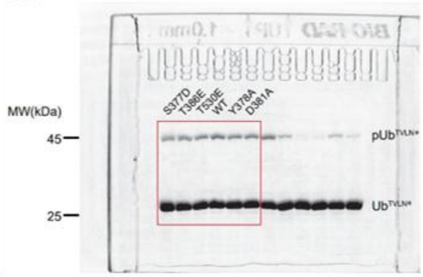


F3-G

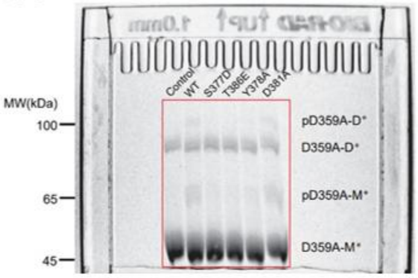


sF3-B

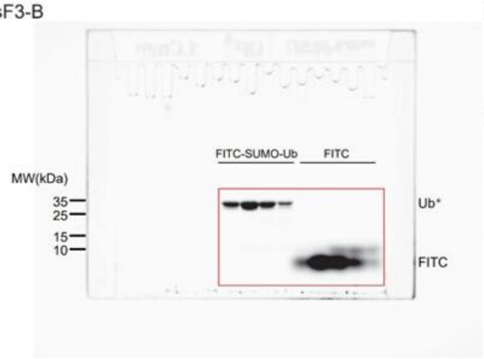


sF3-C

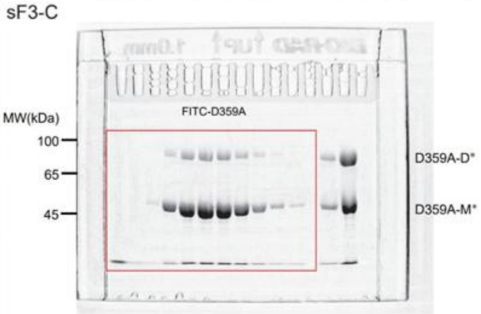


sF3-G

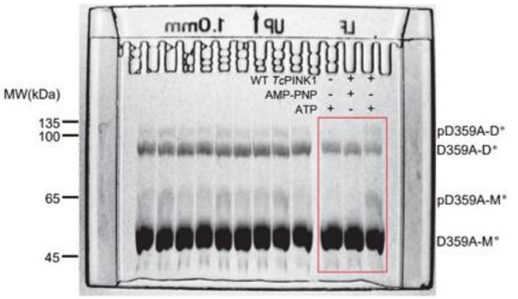


sF3-H

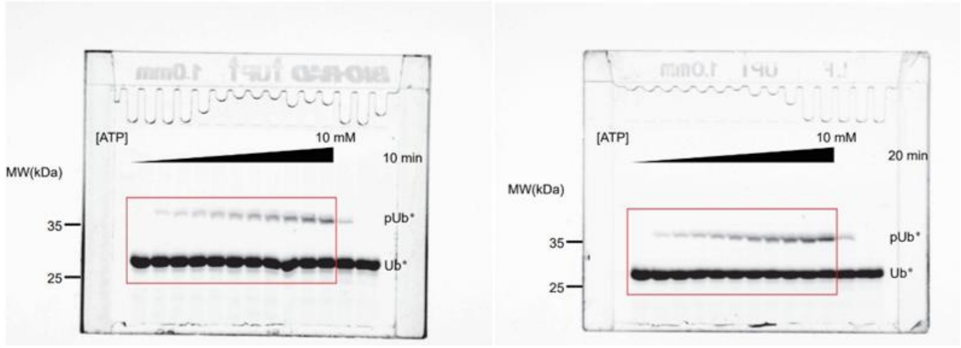

sF4-A

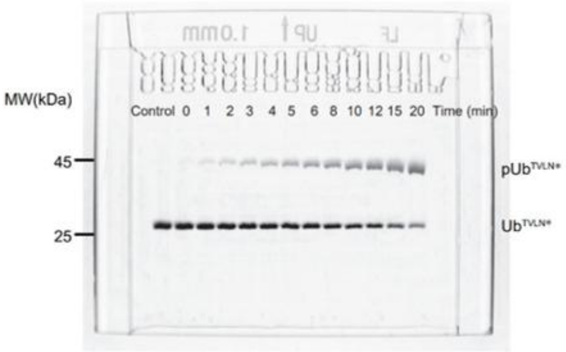


sF4-B

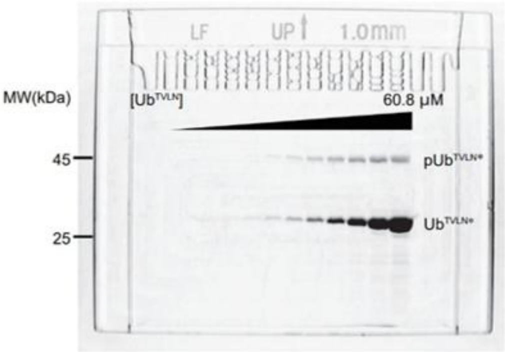


sF4-C

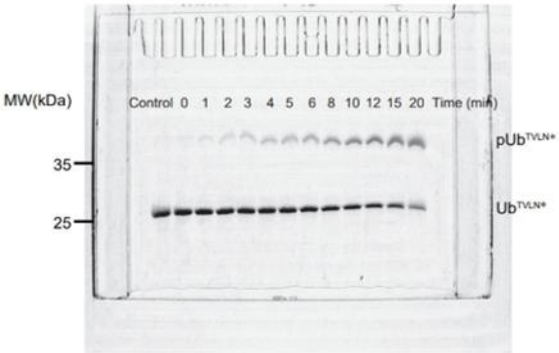


sF4-D

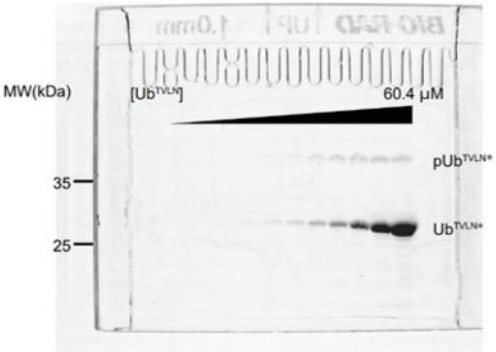


sF4-E

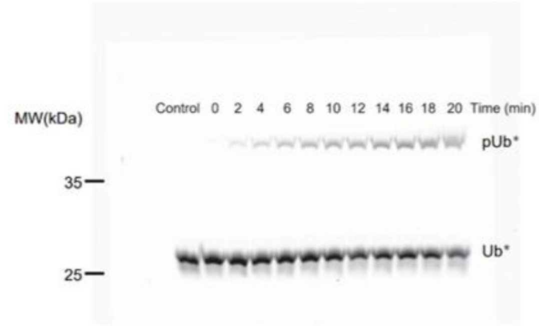


sF4-F

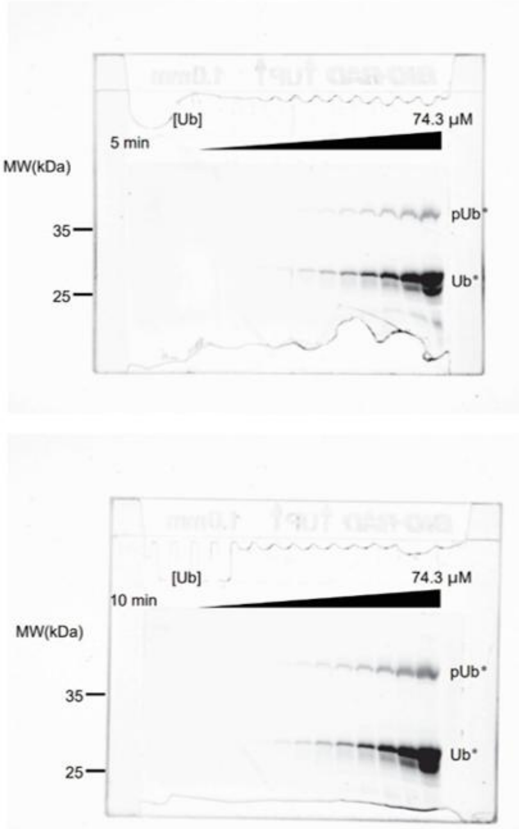

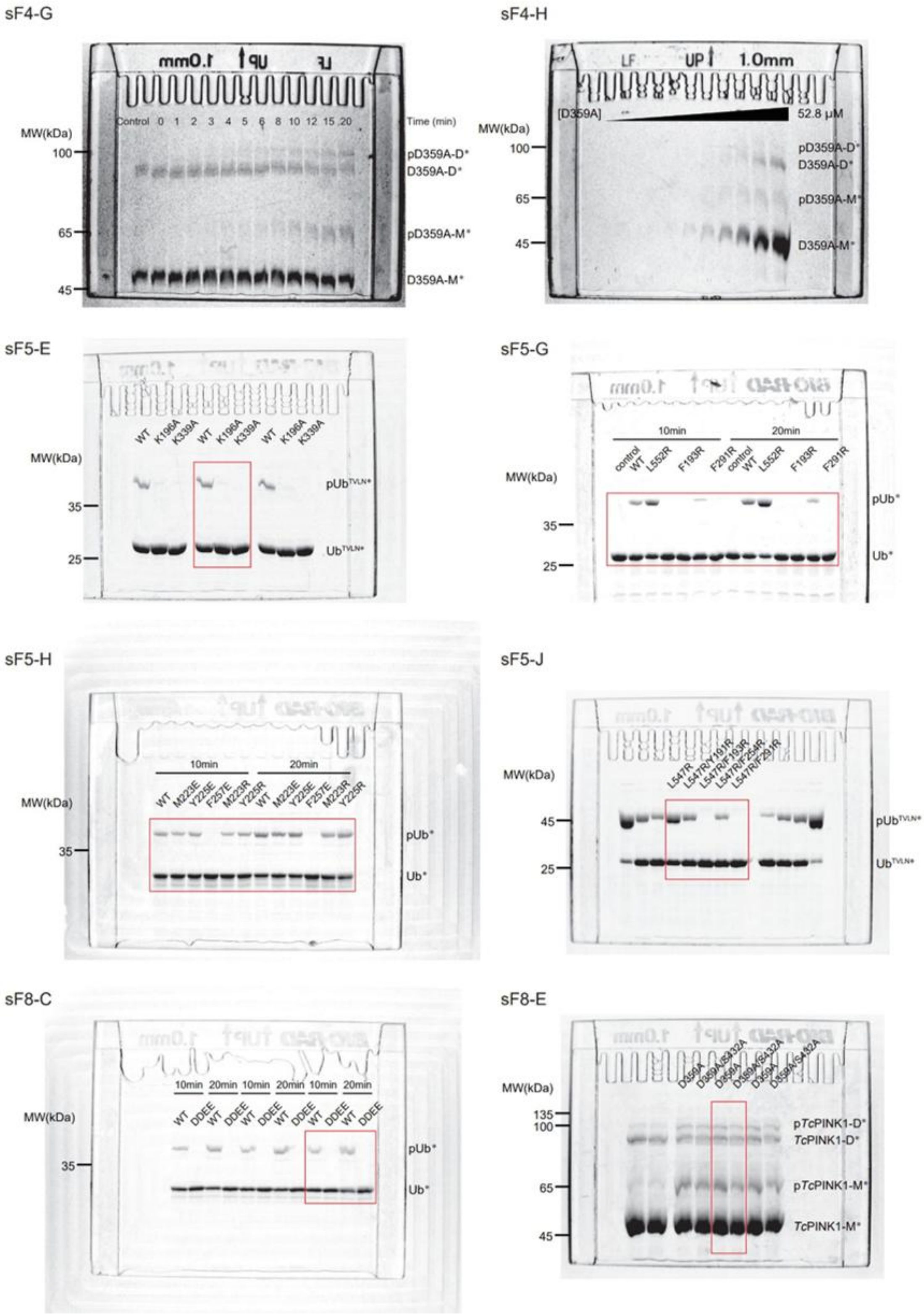


**Fig. S12. Uncropped images of Phos-tag SDS-PAGE gels.**

**Table S1. Cryo-EM data collection, refinement and validation statistics**

| | PINK1$^{117\text{-}570}$ active dimer (EMDB: 80258) (PDB: 25OP) | PINK1$^{L552R}$ symmetric dimer (EMDB: 66272) (PDB: 9WUV) | PINK1$^{L552R}$ (EMDB: 66275) (PDB: 9WUY) | PINK1$^{L552R}$ autoinhibited dimer (EMDB: 66271) (PDB: 9WUU) | PINK1$^{L552R}$ primed activation (EMDB: 66270) (PDB: 9WUT) | PINK1$^{DDEE}$ phosphomimetic dimer (EMDB: 66273) (PDB: 9WUW) | PINK1$^{DDEE}$ tetramer (EMDB: 81366) (PDB: 27RY) |
|---|---|---|---|---|---|---|---|
| **Data collection and processing** | | | | | | | |
| Microscope | FEI Titan Krios | FEI Titan Krios | FEI Titan Krios | FEI Titan Krios | FEI Titan Krios | FEI Titan Krios | FEI Titan Krios |
| Voltage (kV) | 300 | 300 | 300 | 300 | 300 | 300 | 300 |
| Detector | Gatan K3 Summit | Gatan K3 Summit | Gatan K3 Summit | Gatan K3 Summit | Gatan K3 Summit | Gatan K3 Summit | Gatan K3 Summit |
| Magnification | 130,000 × | 130,000 × | 130,000 × | 130,000 × | 130,000 × | 130,000 × | 130,000 × |
| Pixel size (Å) | 0.668 | 0.668 | 0.668 | 0.668 | 0.668 | 0.668 | 0.668 |
| Electron exposure ($e^-$/Å$^2$) | 50 | 50 | 50 | 50 | 50 | 50 | 50 |
| Defocus range (μm) | -1.2 to -1.8 | -1.2 to -1.8 | -1.2 to -1.8 | -1.2 to -1.8 | -1.2 to -1.8 | -1.2 to -1.8 | -1.2 to -1.8 |
| Automation software | EPU | EPU | EPU | EPU | EPU | EPU | EPU |
| Energy filter slit width (eV) | 20 | 20 | 20 | 20 | 20 | 20 | 20 |
| Micrographs used (no.) | 19,212 | 8,201 | 8,201 | 8,201 | 8,201 | 6,127 | 6,127 |
| Symmetry imposed | C2 | C2 | C1 | C1 | C1 | C2 | C1 |
| Initial particle images (no.) | 4,995,315 | 1,861,310 | 1,861,310 | 1,861,310 | 1,861,310 | 1,570,982 | 2,409,694 |
| Final particle images (no.) | 356,219 | 554,347 | 75,286 | 71,484 | 407,577 | 260,669 | 274,915 |
| Map resolution (Å) | 3.48 | 2.81 | 3.22 | 3.35 | 3.04 | 2.51 | 3.46 |
| FSC threshold | 0.143 | 0.143 | 0.143 | 0.143 | 0.143 | 0.143 | 0.143 |
| **Refinement** | | | | | | | |
| Initial model used | 7MP8 | 5YJ9 | 5YJ9 | 5YJ9 | 5YJ9 | 5YJ9 | 5YJ9 |
| Model resolution (Å) | 3.5 | 2.8 | 3.2 | 3.4 | 3.0 | 2.5 | 3.2 |
| FSC threshold | 0.143 | 0.143 | 0.143 | 0.143 | 0.143 | 0.143 | 0.143 |
| Map sharpening *B* factor (Å$^2$) | -177.3 | -147.1 | -115.0 | -129.8 | -150.3 | -119.4 | -117.1 |
| **Model composition** | | | | | | | |
| Non-hydrogen atoms | 6947 | 5626 | 5690 | 5623 | 5587 | 5614 | 11136 |
| Protein residues | 888 | 700 | 711 | 703 | 698 | 702 | 1391 |
| Ligands | 0 | 2 | 1 | 1 | 1 | 0 | 0 |
| *B* factors (Å$^2$) | | | | | | | |
| Protein | 108.35 | 30.11 | 42.27 | 59.28 | 41.41 | 34.91 | 45.61 |

| | | | | | | | |
|---|---|---|---|---|---|---|---|
| Ligand | -- | 39.71 | 34.92 | 45.53 | 57.38 | -- | |
| **R.m.s.deviations** | | | | | | | |
| Bond lengths (Å) | 0.003 | 0.002 | 0.002 | 0.003 | 0.002 | 0.002 | 0.003 |
| Bond angles (°) | 0.652 | 0.515 | 0.490 | 0.509 | 0.496 | 0.490 | 0.603 |
| **Validation** | | | | | | | |
| Molprobity score | 1.93 | 1.36 | 1.41 | 1.68 | 1.48 | 1.31 | 1.78 |
| Clash score | 7.9 | 5.32 | 4.91 | 6.66 | 4.46 | 2.67 | 11.15 |
| Rotamers outliers (%) | 0.26 | 0.00 | 0.00 | 0.32 | 0.00 | 0.00 | 0.00 |
| **Ramachandran plot** | | | | | | | |
| Favored (%) | 91.7 | 97.65 | 97.12 | 95.47 | 96.20 | 96.21 | 96.65 |
| Allowed (%) | 7.73 | 2.05 | 2.88 | 4.39 | 3.80 | 3.79 | 3.35 |
| Outliers (%) | 0.57 | 0.29 | 0.00 | 0.15 | 0.00 | 0.00 | 0.00 |
| **Total Buried surface area at the interface (Å$^2$)** | | 2427 | 2712 | 2735 | 2712 | 3523 | 908 |

**Table S2. Enzyme kinetic parameters of WT *Tc*PINK1 and its mutants**

| Enzyme | Substrate | $K_m$ (×10$^{-4}$ M) | $k_{cat}$ (×10 min$^{-1}$) | $k_{cat}$ /$K_m$ (μM$^{-1}$·min$^{-1}$) |
|---|---|---|---|---|
| *Tc*PINK1$^{117\text{-}570}$ | Ub$^{TVLN}$ | 0.34 ± 0.05 | 17 ± 1 | 4.9 ± 0.3 |
| *Tc*PINK1$^{135\text{-}570}$ | Ub$^{TVLN}$ | 0.09 ± 0.03 | 6.0 ± 0.9 | 7 ± 2 |
| *Tc*PINK1$^{135\text{-}570/L508R}$ Dimer | Ub | 5 ± 1 | 3.3 ± 0.9 | 0.07 ± 0.01 |
| *Tc*PINK1$^{135\text{-}570/L508R}$ Monomer | Ub | 3.1 ± 0.6 | 3 ± 1 | 0.09 ± 0.02 |
| *Tc*PINK1$^{135\text{-}570/L547R}$ | Ub | 3.1 ± 0.3 | 3.2 ± 0.4 | 0.104 ± 0.009 |
| *Tc*PINK1$^{135\text{-}570/L552R}$ | Ub | 4.8 ± 0.3 | 5.3 ± 0.5 | 0.111 ± 0.005 |
| *Tc*PINK1$^{135\text{-}570}$ | Ub | 3 ± 1 | 2.8 ± 0.6 | 0.08 ± 0.01 |
| *Tc*PINK1$^{135\text{-}570}$ | *Tc*PINK1$^{D359A}$ | 0.5 ± 0.1 | 2.18 ± 0.03 | 0.44 ± 0.08 |
| *Tc*PINK1$^{135\text{-}570}$ | ATP | 0.004 ± 0.001 | 0.08 ± 0.01 | 3 ± 2 |

**Movie S1. Animated visualization of the dynamic conformational changes of *Tc*PINK1$^{L552R}$.** The movie displays classified Cryo-EM density maps for 20 conformational states of *Tc*PINK1$^{L552R}$, contoured at 3σ. Low-resolution structural models, refined against these density maps, are represented in cartoon. The animation features 20 consecutive frames of the Cryo-EM density maps, illustrating the dynamic changes in *Tc*PINK1$^{L552R}$. Subunit A and subunit B of the dimeric *Tc*PINK1$^{L552R}$ are colored pink and light blue, respectively. The αC helices are shown in yellow and cyan, while the αi helices are depicted in orange and teal. The autoinhibition loop/β1 is colored green, and α2/β2 is shown in magenta.

**Movie S2. Animated visualization of the dynamic conformational changes of *Tc*PINK1$^{DDEE}$.** The movie features classified Cryo-EM density maps for 20 conformational states of *Tc*PINK1$^{DDEE}$, contoured at 3σ. Low-resolution structural models, refined against these density maps, are depicted in cartoon. The animation shows 20 consecutive frames of the Cryo-EM density maps, illustrating the dynamic changes in *Tc*PINK1$^{DDEE}$. Subunit A and subunit B of the dimeric *Tc*PINK1$^{DDEE}$ are colored pink and light blue, respectively. The αC helices are shown in yellow and cyan; the β1s in orange and blue; the α2/β2s in red and green; and the Ins3s in magenta and violet.
